# Quantitative Profiling of Adaptation to Cyclin E Overproduction

**DOI:** 10.1101/2021.07.26.453751

**Authors:** Juanita C. Limas, Amiee N. Littlejohn, Amy M. House, Katarzyna M. Kedziora, Brandon L. Mouery, Boyang Ma, Dalia Fleifel, Andrea Walens, Maria M. Aleman, Daniel Dominguez, Jeanette Gowen Cook

## Abstract

Cyclin E/CDK2 drives cell cycle progression from G1 to S phase. Cyclin E overproduction is toxic to mammalian cells, although the gene encoding cyclin E (*CCNE1*) is overexpressed in some cancers. To gain insight into how cancer cells tolerate high cyclin E, we extensively characterized non-transformed epithelial cells throughout a time course of chronic cyclin E overproduction. Cells overproducing human cyclin E, but not cyclin D or cyclin A, briefly experienced truncated G1 phases, then endured a transient period of DNA replication origin underlicensing, replication stress, and severely impaired proliferation. Individual cells displayed substantial intercellular heterogeneity in cell cycle dynamics and CDK activity. Each phenotype improved rapidly despite maintaining high cyclin E-associated activity. Transcriptome analysis revealed that adapted cells downregulated a cohort of G1-regulated genes. Withdrawing cyclin E induction only partially reversed the intermediate licensing phenotype of adapted cells indicating that adaptation is at least partly independent of mutations. This study provides evidence that mammalian cyclin E/CDK inhibits origin licensing by an indirect mechanism through premature S phase onset and provides further insight into the relationship between CDK activity and licensing in mammals. It serves as an example of specific oncogene adaptation that may recapitulate molecular changes during tumorigenesis.

## INTRODUCTION

Cell cycle regulation depends on tight control of cyclin protein expression (Evans *et al*., 1983). Cyclins activate cyclin-dependent protein kinase enzymes (CDK) to govern cell cycle progression. In G1 phase, growth factors induce cyclin D expression to activate CDK4 and CDK6. This complex phosphorylates the E2F-inhibitor retinoblastoma protein (RB), relieving repression of E2F, and driving transcription of a suite of genes necessary for S phase entry, including *CCNE1* (DeGregori *et al*., 1995; Dimova and Dyson, 2005; Narasimha *et al*., 2014; Sanidas *et al*., 2019). Rb hyperphosphorylation by cyclin E/CDK2 (and/or cyclin D/CDK4-6) fully inactivates Rb, and E2F is then fully active (Zarkowska and Mittnacht, 1997; Yang *et al*., 2020). Cells can then progress from late G1 into S phase and initiate DNA replication. Importantly, cyclin E is overproduced in many cancers as a consequence of gene amplification or dysregulated transcription (Chu *et al*., 2021), yet high cyclin E can induce premature S phase onset leading to replication stress, proliferation failure, and genome instability (Minella *et al*., 2002; Teixeira *et al*., 2015a). However, the mechanisms of cyclin E-induced replication stress are not yet fully understood.

Premature S phase onset shortens the time necessary for essential molecular processes in G1 phase, and failure to complete these steps may contribute to replication stress in S phase. One of the essential G1 processes is DNA replication origin licensing. In mammalian cells, thousands of chromosomal sites are prepared through the loading of MCM complexes to render them competent for replication initiation in S phase. MCM complexes are loaded to license origins by the concerted action of the Origin Recognition Complex (ORC), and the CDC6 and CDT1 proteins (Gillespie *et al*., 2001; Evrin *et al*., 2009; Remus *et al*., 2009). As soon as S phase starts, ORC, CDC6, and CDT1 are inactivated to completely stop further MCM loading to avoid re-licensing and re-replication. Re-replication is a form of endogenous DNA damage and genome instability that can contribute to oncogenesis (Green and Li, 2005; Davidson *et al*., 2006; Arias and Walter, 2007; Zhou *et al*., 2020).

Because licensing is tightly restricted to G1 phase by the inactivation of MCM loading factors outside G1, all of the licensing needed to support complete genome duplication in S phase must occur before the G1/S transition. One of the many challenges during S phase is replication fork stalling which can occur stochastically from DNA lesions, replication fork damage, collisions with transcription complexes, repetitive sequences, or other barriers (Ait Saada *et al*., 2018; Berti *et al*., 2020). Stalled forks can be accommodated by activating (or “firing”) additional nearby origins (Woodward *et al*., 2006). To ensure that there are enough licensed origins available to serve as both primary origins and in a rescue capacity, cells load excess MCM during G1 to license many dormant origins (Ge *et al*., 2007). Without dormant origins, cells are hypersensitive to replication inhibitors, exhibit higher genome instability, and are more prone to tumor formation (Pruitt *et al*., 2007; Shima *et al*., 2007; Ibarra *et al*., 2008; Kawabata *et al*., 2011).

We and others had previously shown that cyclin E overproduction in non-transformed cells not only induces replication stress and DNA damage, but also shortens G1 phase (Resnitzky *et al*., 1994; Matson *et al*., 2017). Moreover, persistent cyclin E expression in non-transformed cells is toxic over a period of days or weeks (Minella *et al*., 2002; Jones *et al*., 2013; Kok *et al*., 2020). We were thus intrigued by the fact that some tumor-derived cell lines with high cyclin E proliferate well (Barretina *et al*., 2012; Asghar *et al*., 2017; Geng *et al*., 2018). In addition to shortening G1 phase, ectopic cyclin E overproduction induces cells to enter S phase with substantially less loaded MCM than control cells (Matson *et al*., 2017). These observations are consistent with the short-term origin licensing inhibition from transient cyclin E overproduction even in cancer cells that already express high cyclin E (Ekholm-Reed *et al*., 2004) and a general understanding that CDK activity directly inhibits MCM loading factors (Nguyen *et al*., 2001; Diffley, 2004; Zielke *et al*., 2011; Zielke *et al*., 2013). If cyclin E is toxic to cells, then how do cancer cells tolerate cyclin E overexpression and reduced origin licensing over the long timelines required for tumor development? Such cells presumably have mechanisms to accommodate the cell cycle and replication stress effects of high cyclin E, but those mechanisms are still unknown.

To investigate potential mechanisms to tolerate cyclin E overexpression, we have extensively analyzed multiple independent time courses throughout the process of adapting to ectopic cyclin E overproduction in non-transformed epithelial cells. Using non-transformed cells in this study allows us to avoid the confounding variables of genetic and epigenetic alterations present in cancer cells to better understand the effects of cyclin E overproduction. We characterized the acute effects of cyclin E overproduction at single cell resolution and analyzed cell populations as they adapted to this particular stress. We found that high cyclin E, but not high cyclin D or cyclin A, initially induces cell cycle changes, defects in origin licensing, and replication stress, and we provide evidence that the licensing defects are caused more by truncating G1 than by direct MCM loading factor inactivation. Cells then adapt over a period of just a few weeks to lengthen G1 phase to improve origin licensing while maintaining cyclin E expression and activity. We extensively quantified the cell cycle, replication, and gene expression changes during this cellular adaptation as a potential model for how cancers tolerate cyclin E overproduction.

## RESULTS

### Cyclin E overproduction shortens G1

To explore the specific cellular consequences of cyclin E overproduction we generated non-transformed human retinal pigmented epithelial cell lines with inducible cyclin E, cyclin D, or cyclin A cDNAs. We included cyclin D for comparison because it is also overproduced in some cancers but is not associated with changes in origin licensing and replication stress, and we included cyclin A because it is an alternative activator of CDK2. We established cell lines with one isoform of each cyclin (cyclin D1, cyclin E1, and cyclin A2) and isolated colonies from single cells to generate monoclonal populations. We first treated cells with varying concentrations of dox for 48 hours (~two cell cycles) and probed whole cell lysates by immunoblotting to measure cyclin overproduction (**Figure 1A**). Each cyclin was overproduced in a dose-dependent manner (**Figure 1B**). We were particularly interested in cyclin E overproduction levels relative to those that may be achieved in tumors. We thus examined mRNA overexpression of *CCNE1* in The Cancer Genome Atlas (TCGA) and noted similar increases in some breast cancer tumors compared to normal tissue. Both cyclin E and cyclin A are products of E2F-regulated genes (Ohtani *et al*., 1995; Schultze *et al*., 1995) and are thus repressed by Rb. Because G1-S CDK activity de-represses Rb, we also tested for indirect effects of overproducing one cyclin on the others (Chellappan *et al*., 1991; Burkhart and Sage, 2008; Rubin *et al*., 2020). Immunoblotting for each G1-S cyclin showed little to no consistent effect of overproducing one cyclin on expression of another cyclin with the possible exception of cyclin A inducing cyclin E (**Figure 1A**). We show darker exposures in Figure 1A to make endogenous cyclins visible, but none of these signals were outside the linear detection range, and we include lighter exposures in **Supplemental Figure S6**.

**Figure 1.**
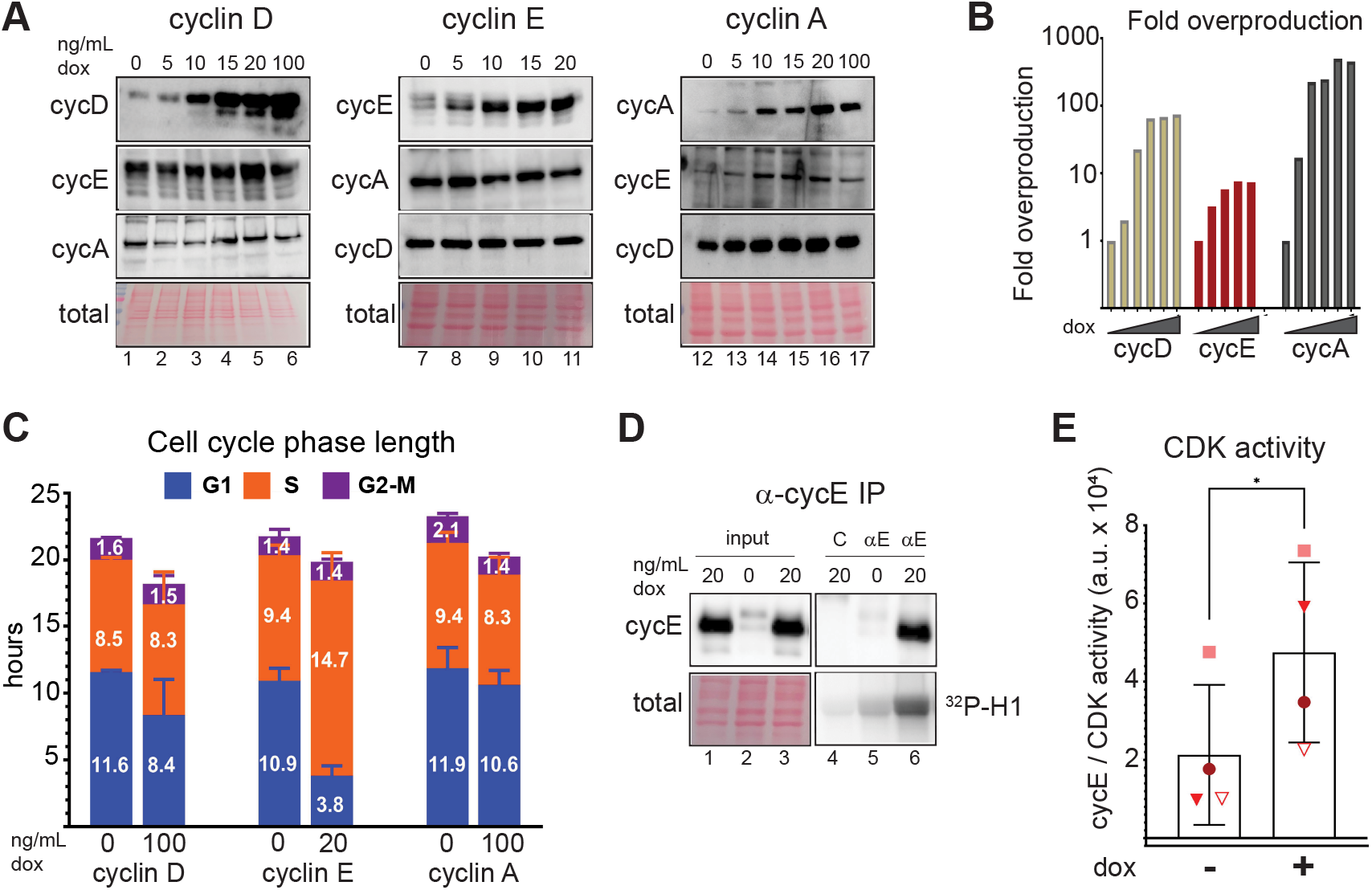
Cyclin E overproduction shortens G1. (A) Non-transformed retinal pigmented epithelial (RPE1-hTert) cells with stably-integrated doxycycline-inducible human cyclin D1 (left), cyclin E1 (middle), or cyclin A2 (right) were treated with the indicated concentrations of doxycycline (dox) for 48 hours prior to immunoblotting for the indicated ectopic and endogenous proteins (representative of 3 replicates). (B) Fold overproduction of each induced cyclin relative to endogenous levels from A. (C) Cell cycle phase lengths and distributions determined by flow cytometry and doubling time after 48 hours of cyclin induction with 100 ng/ml (cyclin D1), 20 ng/ml (cyclin E1), and 100 ng/ml (cyclin A2) doxycycline (n=3). Difference from control in G1 length; cyclin D *, cyclin E ****, cyclin A not significant. (D) Cyclin E and histone H1 kinase activity in immunoprecipitates of cyclin E from lysates measured after 48 hours of cyclin E overproduction (20 ng/mL). The control immunoprecipitate “C” is lysate from induced cells treated with non-immune serum. (E) Quantification of four independent replicates of (D). *p ≤ 0.05, **p ≤ 0.005, ***p ≤ 0.0005.

We selected one doxycycline dose for each cell line for further study based on the maximum dose that did not induce an immediate proliferation arrest. These concentrations were 100 ng/ml for cyclin D and cyclin A but 20 ng/ml for cyclin E; 100 ng/ml induced strong proliferation arrest in the cyclin E line (not shown). We then tested for changes in cell cycle phase lengths by combining the doubling time of each cell line with the percentage of cells in G1, S, or G2/M using flow cytometry analysis of both DNA content and DNA synthesis (EdU incorporation) (**Supplemental Figure S1A-B**). We observed no evidence of cell death (e.g., floating cells) in response to cyclin overproduction. Overproducing cyclin E dramatically shortened G1 by nearly three-fold (adjusted p-value <0.0001) whereas overproducing cyclin D or cyclin A minimally shortened G1 (**Figure 1C**). To directly evaluate the consequence of cyclin E overproduction on the activity of its kinase partners, we also quantified CDK / cyclin E-associated histone H1 kinase activity in cyclin E immunoprecipitates from lysates of asynchronously proliferating cells and quantified an average two-fold increase in CDK activity (**Figure 1D**, N=4 replicates quantified in **Figure 1E**). The differences between fold-induction of cyclin E protein and fold-activation of CDK activity could reflect limiting CDK2 and/or compensatory mechanisms to restrain cyclin E/CDK2 activity.

### Cyclin E overproduction causes underlicensing and replication stress

The short G1 phase in cyclin E-overproducing cells raised the possibility that cells entered S phase with less than the normal amount of loaded MCM to license origins. To test this idea, we analyzed the amount of loaded MCM per cell in cells overproducing cyclin E in early S phase using an analytical flow cytometry approach (**Figure 2A**). Because MCM is unloaded over the course of S phase, we focused only on early S phase cells as a measure of the maximum amount of MCM that had been loaded in the previous G1 phase. Briefly, we pulse-labeled asynchronously proliferating cells with EdU prior to harvesting to identify S phase cells. We then treated cells with non-ionic detergent to remove soluble MCM leaving behind chromatin-bound MCM. We fixed the extracted cells, immunostained for endogenous MCM2 (as a marker of the MCM2-7 complex) and stained for DNA content with DAPI as described in Materials & Methods. Flow cytometry data are shown as bound MCM vs. DNA content with color coding as follows: MCM signal below antibody threshold is indicated in grey, and cells with detectable bound MCM but no EdU incorporation are indicated in blue. EdU positive cells (S phase) are indicated in orange (**Figure 2B**). The amount of loaded MCM per cell in very early S phase indicates the abundance of licensed origins available for the duration of that S phase; we identified this early S phase subpopulation in green (**Figure 2C** and **Supplemental Figure S1A** and **B**). Cyclin E overproduction reduced the amount of MCM that had been loaded at the time of S phase entry (**Figure 2C**, green). In contrast, cyclin D or cyclin A overproduction only modestly reduced MCM loading in early S phase (**Supplemental Figure S1C-F**). To compare control and treated cell populations, we plotted the MCM intensities in just the early S phase cells from **Figure 2C** on a single histogram. The shift of MCM loading distribution to the left in cyclin Eoverproducing cells indicates S phase entry with less origin licensing (**Figure 2D**). Similar histograms of early S MCM loading in cells overproducing cyclin D and cyclin A indicate no significant effects on licensing in these populations (**Figure 2D**). We also plotted a combination of 20 biological replicates showing single cell MCM signal intensities on one graph (**Figure 2E**, raw intensities in arbitrary units). We observed differences in the magnitude of underlicensing among independent replicates, but cyclin E overproduction consistently induced significant underlicensing. Across many experiments, cyclin E overproduction reduced licensing in early S by 2.5-fold relative to control cells.

**Figure 2.**
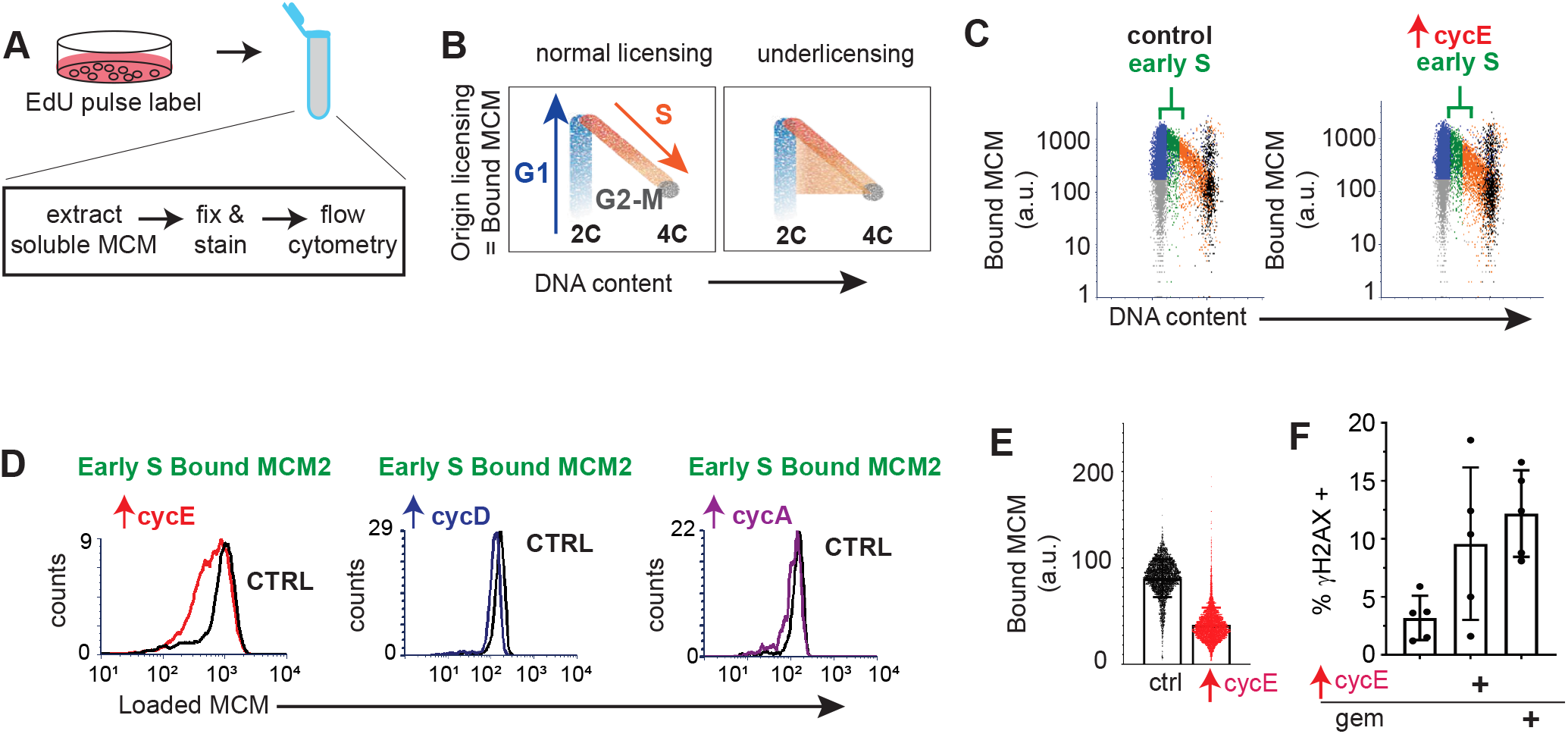
Cyclin E overproduction causes underlicensing and replication stress. (A) Workflow for quantifying chromatin-bound endogenous MCM by flow cytometry. (B) Illustration of dot plots of DNA content (x-axis), and bound MCM2 as a marker of the MCM2-7 complex (y-axis). Individual cells are colored blue for G1 MCM-positive, orange for EdU-positive and MCM-positive (S phase), and gray for MCM-negative. Typical patterns for normal licensed (left) and underlicensed (right) populations. (C) Analysis of MCM loading in control and cyclin E-overproducing cells after 48 hours of induction with 20 ng/mL dox. Early S phase cells (defined as MCM-positive, EdU-positive, G1 DNA content) are plotted in green and indicated with brackets (representative of a minimum 6 replicates). (D) Loaded MCM in early S phase cells plotted as intensity per cell in response to overproducing cyclin E (left, same cells as in C), cyclin D (middle) and cyclin A (right). (E) Early S phase MCM loading intensity in single cells from a total of 20 independent replicates. (F) Cells were treated with 20 ng/ml dox to induce cyclin E overproduction or 1 nM gemcitabine for 48 hours as indicated then analyzed by flow cytometry for % γ-H2AX positive cells. Positive cells were defined in Supplemental Figure S2 (n=5).

Cells overproducing cyclin E for 2-3 days were still proliferating despite reduced origin licensing. We reasoned that these cells had licensed enough primary origins in G1 to usually complete S phase, but there would be too few licensed dormant origins to suppress replication stress and genome instability. Previous work by us and many others established that substantially reducing origin licensing leads to accumulation of DNA damage markers, such as γ-H2AX, and impaired proliferative fitness (Orr *et al*., 2010; Alvarez *et al*., 2015; Bai *et al*., 2016) induces replication stress. To determine if these cells were functionally “underlicensed,” we quantified γ-H2AX as a marker of replication stress (and also conducted immunostaining for 53BP1 nuclear bodies in **Figure 4A**.) (Chanoux *et al*., 2009; Zeman and Cimprich, 2014). We quantified the percentage of γ-H2AX-positive cells using analytical flow cytometry in cells overproducing cyclin E for 48 hrs compared to untreated cells or cells exposed to the replication inhibitor gemcitabine as a positive control for replication stress. Cells overproducing cyclin E for just 48 hours generated more γ-H2AX positive cells than control cells (**Figure 2F**, gating strategy for scoring γ-H2AX provided in **Supplemental Figure S2**). We thus conclude that cyclin E overproduction generates functionally underlicensed cells that experience replication stress.

### Proliferation and licensing perturbations during adaptation to chronic cyclin E overproduction

We were curious if cyclin E-overproducing cells could sustain long-term proliferation while severely underlicensed. We therefore monitored cell proliferation during more than a month of continuous induction for each of the cyclin-overproducing lines by plotting days in culture on the x-axis and the inverse of the population doubling time in hours on the y-axis (1/Dt, e.g. 20 hr doubling time is 0.05 on the y-axis). In the first 3 days, each of the three G1-S cyclins accelerated proliferation, but only cyclin D sustained consistent accelerated proliferation whereas cells overproducing cyclin A showed only modest acceleration relative to controls (**Supplemental Figure S3A**). In contrast, cyclin E-overproducing cells rapidly slowed after the initial burst of faster proliferation (**Figure 3A**). A prior study documented strong proliferation defects after two weeks of cyclin E overproduction in human fibroblasts (Minella *et al*., 2002), and we made similar observations in these epithelial cell lines; cultures grew much more slowly after 2-3 weeks of cyclin E overproduction. Surprisingly, the populations recovered after this period of slow proliferation and returned to accelerated growth by week 4-5. Normal doubling time of these cells is 22-23 hours, whereas cells overproducing cyclin E had doubling times that ranged from 15 hours during the “acute” phase in the first few days to 81 hours during the “proliferative crisis” after 2-3 weeks. By the end of the experiments all cyclin-overproducing cultures were proliferating somewhat faster than controls (**Figure 3A** and **Supplemental Figure S3A**). Interestingly, the exact timing of the proliferative crisis varied within a ~two-week window, but all populations were proliferating well by day 33. We did not observe additional proliferation rate changes after day 30 (not shown).

**Figure 3.**
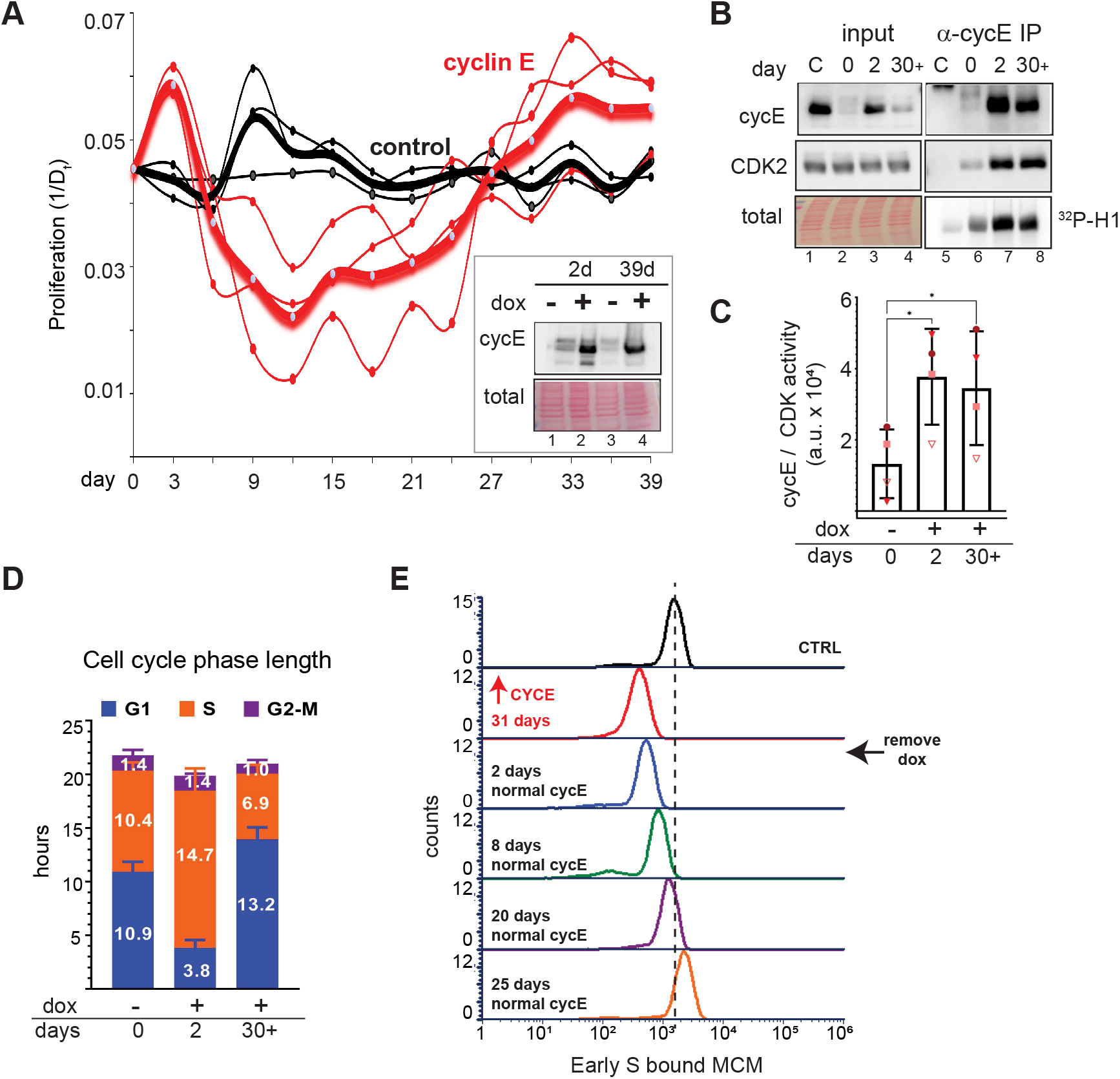
Significant proliferation and licensing perturbations from adaptation to chronic cyclin E overproduction. (A) Proliferation rate measured every three days in control cells and cells overproducing cyclin E; heavy lines are the means and thin lines are 3 independent replicates. Doubling time in hours (Dt) was calculated, and proliferation is plotted as the inverse of the population doubling time in hours on the y-axis (1/Dt) and days in culture on the x-axis. Inset: Representative immunoblot analysis of cyclin E in cells grown for 2 days or for more than 30 (39) days in 20 ng/ml dox (lanes 2 and 4). Lane 3 is cyclin E expression in adapted cells 48 hours after withdrawing dox. (B) Cyclin E, endogenous CDK2, and histone H1 kinase activity in immunoprecipitates of cyclin E from lysates prepared after 2 or 30+ days of proliferation in 20 ng/mL dox. C= control serum (C) Kinase activity in four independent replicates of B. (D) Cell cycle phase lengths after 2 or 30+ days of proliferation in 20 ng/mL dox (n= 3 replicates). Difference from control in G1 length: control vs acute p ≤ 0.00005, acute vs 30+ days \*\**p* ≤ 0.005. (E) Early S MCM loading in control cells, adapted cells at day 31, or cells at the indicated days of culture after removing dox (representative of 3 replicates). Dotted line represents midpoint of the control histogram as a reference for other samples. No additional differences in phenotype were apparent after day 25.

We were intrigued by the pattern of slowed proliferation, or “proliferative crisis,” then recovery in the cyclin E-overproducing cells and sought to characterize the adapted populations in more detail. We first analyzed the levels of induced cyclin E protein in cells after 39 days of chronic cyclin E expression compared to cyclin E levels after just two days. Interestingly, adapted populations still overproduced cyclin E (**Figure 3A inset**, compare lanes 2 and 4) and when dox was removed for 48 hours, cyclin E expression reverted back to endogenous levels (**Figure 3A inset**, compare lanes 3 and 4). We obtained similar results for cyclins D and A (**Supplemental Figure S3B**, compare lanes 2 & 4 and lanes 6 & 8). We analyzed several independently-derived cyclin E-adapted populations, and sometimes found reduced cyclin E protein in adapted cells compared to the initial levels after induction (for example, **Figure 3B**, compare lanes 3 and 4), but the degree of protein downregulation varied among replicates. In all replicates however, cyclin E levels were still higher in adapted populations than endogenous cyclin E. Importantly, the amount of CDK <u>activity</u> associated with cyclin E overproduction remained high in adapted cells similar to cells shortly after cyclin E induction (**Figure 3B**, compare lanes 7 & 8 and **Figure 3C**). In addition, CDK2 protein in cyclin E immunoprecipitates was also similar (**Figure 3B**, compare lanes 7 and 8). We then analyzed the cell cycle phase lengths in adapted populations relative to control cells or cells that overproduced cyclin E for only 2 days. We once again observed the extreme shortening of G1 phase as an acute early response to cyclin E overproduction (mean 10.9 hours in control vs 3.8 hours after 2 day), but strikingly, despite their elevated cyclin E-associated activity, adapted cells spent at least as much time in G1 as control cells did (**Figure 3D**, 10.9 hours in control vs 13.2 hours after 30 or more days).

One mechanism by which high cyclin E could induce underlicensing is by shortening G1 phase without increasing the rate of MCM loading in G1. In this scenario, cells start S phase before the normal amount of MCM has been loaded. We hypothesized that if this premature S phase is the primary mechanism causing underlicensing, then adapted cells with longer G1 phases would start S phase less underlicensed than cells after acute cyclin E overproduction. We thus analyzed MCM loading in adapted cells compared to uninduced cells. Interestingly, the adapted cells were not as severely underlicensed as cells with very short G1 phases (Figure 2) was also not as high as in uninduced cells (**Figure 3E**, compare control to ↟CycE 31 days). This intermediate level of licensing was clearly compatible with robust cell proliferation despite the high cyclin E-associated kinase activity.

To test if the intermediate licensing was solely dependent on elevated cyclin E, we monitored licensing in adapted cells after withdrawing doxycycline. Removing doxycycline from adapted cells resulted in a rapid return to endogenous cyclin E levels (**Figure 3A inset**, compare lanes 3 and 4). Cells proliferated normally and did not show evidence of experiencing a second proliferative crisis which indicated that adaptation did not include acquiring dependence on high cyclin E (not shown). Surprisingly, returning cyclin E to endogenous levels after 30 days of adaptation to high cyclin E did not immediately rescue the intermediate licensing of adapted cells. We analyzed cells at 2, 8, 20, and 25 days after doxycycline withdrawal and quantified a gradual improvement in origin licensing over the course of this recovery period (**Figure 3E**, compare adapted cells after 31 days of dox induction in red to days 20 and 25 after dox removal in purple and orange). Cells proliferated normally during the recovery and showed no evidence of experiencing additional proliferative stress (not shown). The finding that the licensing phenotype in adapted cells did not quickly revert to the parental phenotype as soon as cyclin E was no longer overproduced suggested that adaptation is associated with persistent physiological changes which are independent of cyclin E but that reverse after many cell divisions with normal cyclin E levels. Furthermore, the fact that cells eventually returned to normal licensing indicated that adaptation was not solely through selection for permanent genetic changes (although mutations may have occurred). In further support of non-genetic adaptation, we did not observe rare clones arising from a field of mostly arrested cells during the proliferative crisis.

### Chronic cyclin E overproduction induces transient DNA damage response markers and heterogeneous cell cycle arrest

We next sought to characterize molecular markers that may have changed throughout adaptation. We anticipated that cells accumulate endogenous DNA damage during the first weeks as a consequence of chronic underlicensing and replication stress. To test this notion, we fixed cells on coverslips every 3 days during chronic cyclin E overproduction and stained for the replication stress and DNA damage marker, 53BP1 (Schultz *et al*., 2000). We analyzed a single population of cells going through adaptation to cyclin E overproduction for 33 days by quantifying the number of 53BP1 nuclear bodies in thousands of cells from 20 fields of view in each timepoint (**Supplemental Figure S4A**). We chose a conservative arbitrary threshold for automated counting to quantify the brightest 53BP1 nuclear bodies (see Methods). We classified cells as having one bright nuclear body, multiple nuclear bodies, or no nuclear bodies. Prior to induction, cells produce some 53BP1 nuclear bodies as a result of constitutive replication stress in unperturbed cells (Moreno *et al*., 2016). Within the first few days of cyclin E overproduction, we found consistently more cells with one or more 53BP1 nuclear bodies, and this increase spanned the first two weeks of chronic cyclin E overproduction and then subsided back to control levels (**Figure 4A**). We note that the period of elevated 53BP1 nuclear bodies generally coincided with the time of proliferative crisis and the return to baseline 53BP1 signals coincided with the return to accelerated proliferation in **Figure 3A**.

**Figure 4.**
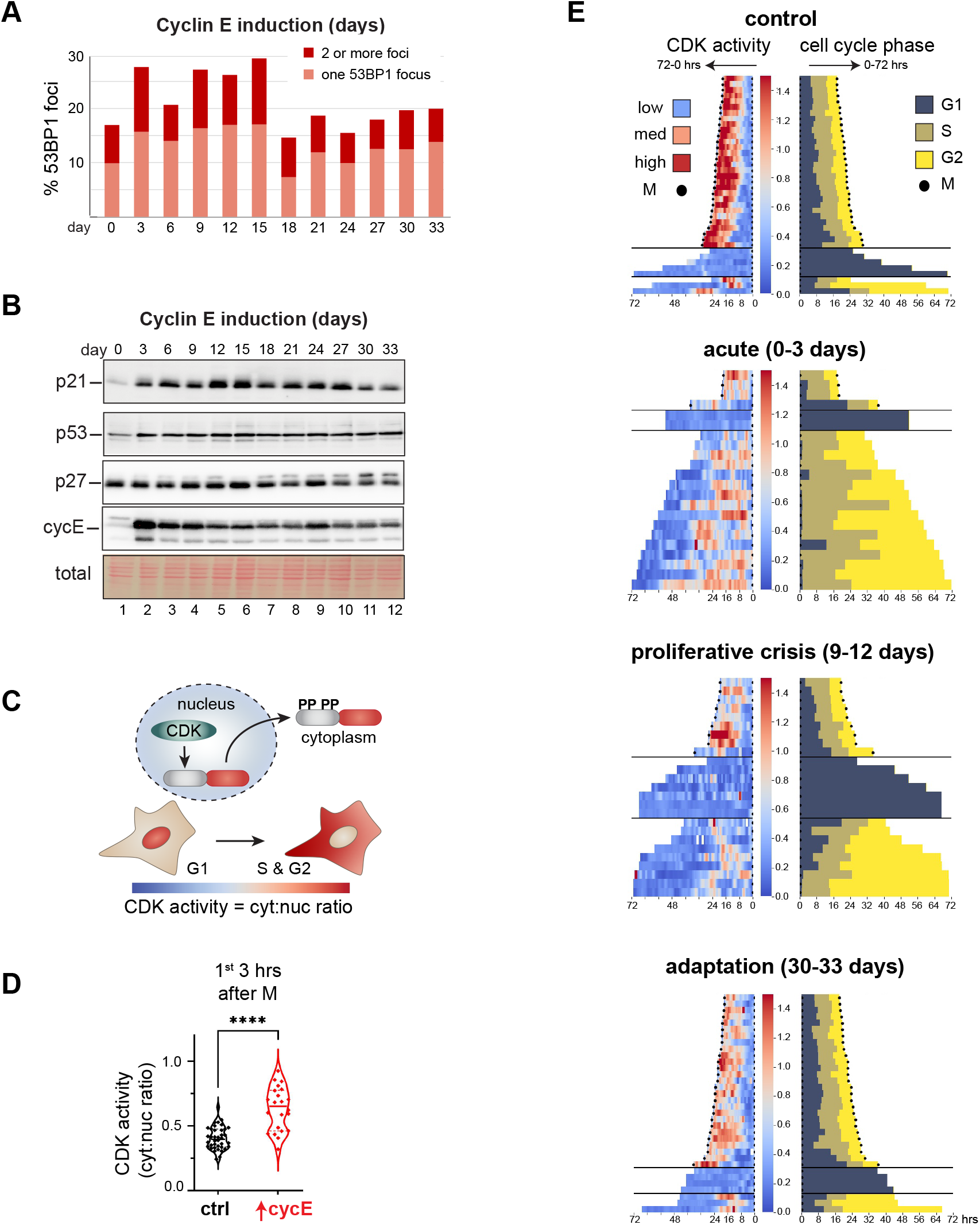
Chronic cyclin E overproduction induces transient DNA damage response markers and heterogeneous cell cycle arrest. (A) Cells were induced with 20 ng/mL doxycycline to overproduce cyclin E and stained for endogenous 53BP1 nuclear bodies at the indicated time points; cells were classified as having 1 or more than one bright 53BP1 nuclear body. (B) Endogenous p21, p27 and p53 plus both ectopic and endogenous cyclin E in lysates of cells treated as in (A). (C) Illustration of the translocation-based CDK1/CDK2 activity biosensor from Spencer et al., 2013. (D) CDK activity in the first three hours of G1 after cyclin E induction plotted as the cytoplasmic to nuclear ratio of biosensor localization (n=60 cells\*\*\*\**p* ≤ 0.00005). (E) CDK activity (left) and cell cycle phase lengths defined by PCNA localization (right) during 72 hours of live imaging at the indicated times after continuous cyclin E overproduction. Individual cell traces begin with cell division and extend left for CDK activity and right for cell cycle phases. CDK activity is plotted from low (blue) to high (red) cytoplasmic:nuclear biosensor localization. Black dots indicate mitosis.

The increase in 53BP1 nuclear bodies indicated DNA damage that may have triggered a cell cycle checkpoint and contributed to the proliferative crisis. We therefore probed lysates of cells experiencing chronic cyclin E overproduction collected every 3 days for the induction of p53 and its downstream target, p21, both well-established markers of the DNA damage response (El-Deiry *et al*., 1993; Macleod *et al*., 1995; Kubbutat *et al*., 1997). We detected a minor and sustained elevation in p53 protein levels throughout most of the time course, but a marked induction of p21 (**Figure 4B**). The induction of the p21 CDK inhibitor preceded the proliferative crisis and lasted past the return to rapid proliferation in this example. In contrast, the p27 CDK inhibitor was unaffected. The p21 protein is degraded in S phase cells, but p21 is stable in G1 cells (Abbas *et al*., 2008; Nishitani *et al*., 2008). Given that adapted cells spend more time in G1 and less time in S phase (**Figure 3D**), the elevated p21 levels in lysates in later time points could reflect cell cycle phase distribution, a persistent DNA damage response, or both. On the other hand, p21 induction at earlier time points is more likely to be attributable to DNA damage since G1 phases were much shorter, and 53BP1 nuclear bodies were more abundant at those times. It is possible that this early p21 induction limited the amount of kinase activity the overproduced cyclin E could stimulate (**Figures 1E** and **3C**). We also detected slightly elevated p21 induction from the addition of neocarzinostatin (NCS), a DNA damage-inducing agent, in cells after early cyclin E overproduction compared to adapted cells (**Supplementary Figure S4B**). Our working model is that adaptation includes a long, but transient activation of the DNA damage response.

Independent adaptations varied in the dynamics of the proliferation responses (**Figure 3A**). Nevertheless, the pattern was consistent in each replicate: cultures initially proliferated rapidly, then very slowly, then returned to rapid proliferation. We routinely monitored cultures in proliferative crisis and observed heterogeneous morphologies with some large cells reminiscent of senescence, some cells with elongated shapes, and others that resembled normal proliferating cells (not shown). We were therefore interested in understanding adaptation in single cells. The biochemical kinase activity analysis in **Figures 1 and 3** could not distinguish cyclin E-associated activity in G1 cells from activity in other cell cycle phases. To further examine single cell CDK activity in real time we introduced a single cell reporter that responds to both CDK1 and CDK2 activity by translocating from the nucleus to the cytoplasm upon CDK-mediated phosphorylation (**Figure 4C**). This reporter was generated from a fragment of the DHB CDK substrate, does not respond to CDK4/6, and has been extensively characterized in prior studies (Gu *et al*., 2004; Hahn *et al*., 2009; Spencer *et al*., 2013; Schwarz *et al*., 2018; Yang *et al*., 2020). In addition, we introduced a PCNA-based reporter to mark the boundaries of S phase by changes in localization of PCNA (Leonhardt *et al*., 2000). To monitor CDK1/2 activity, we measured the cytoplasmic-to-nuclear fluorescence intensity ratio of the reporter: a low ratio indicates low CDK activity, while increasing ratios from cytoplasmic reporter translocation indicate increasing CDK activity. We first tested the immediate effects of cyclin E induction by analyzing G1 cells in the first 3 hours after mitosis shortly after doxycycline addition. As expected, high cyclin E induced a strong spike in CDK1/2 activity evidenced by increased reporter translocation and increased cytoplasm-to-nucleus fluorescence (**Figure 4D**).

We then subjected this reporter line to chronic cyclin E overproduction and collected time lapse fluorescence images for 72 hours at four distinct timepoints: (1) control, (2) “acute” cyclin E overproduction (0-3 days), (3) proliferative crisis (9-12 days), and (4) adaptation 30-33 days. For individual cells we calculated CDK activity and cell cycle phase lengths or arrest; we restricted analysis to cells for which the first mitosis was visible and did not include the cells that did not divide during the entire 72 hours. We generated heat maps of CDK1/2 activity on the left and tracks for cell cycle phases on the right, and each trace begins with mitosis and ends with mitosis or the end of imaging (**Figure 4E**). In the control population with no cyclin E overproduction, CDK1/2 activity followed the expected pattern that was low in G1 (blue) and increased through S and G2 phases (pink and red) followed by mitosis (black dots) for most cells.

Although these cultures started from monoclonal populations, we found wide intercellular heterogeneity in CDK1/2 and cell cycle dynamics. At each timepoint we identified three discrete patterns: 1) a complete cell cycle that ended with mitosis, 2) mitosis followed by a G1/G0 arrest, and 3) cells that appeared to complete S phase but never divided. The first three days of cyclin E overproduction (“acute”) were characterized by many cells with extremely short G1 phases (dark blue bars) and long S phases, as expected from the flow cytometry analysis in **Figure 3D**, but also less pronounced oscillations in CDK activity compared to controls. Many of these cells completed S phase but never entered mitosis or underwent nuclear envelope breakdown. Instead, these “G2-arrested” cells returned to low CDK activity as though they had skipped mitosis, but they did not start S phase again during the imaging window (**Figure 4E, acute (0-3 days)**).

In contrast, populations in the proliferative crisis had many more individual cells that divided but then entered very long G1 phases and neither increased CDK activity nor progressed to S phase, and many cells completed S phase but arrested in the subsequent G2 with low CDK activity (**Figure 4E, proliferative crisis (9-12 days)**). Many cells were also entirely non-dividing with no mitoses visible for the entire 72 hours (not shown). Nonetheless, a fraction of cells completed a full cell cycle although they displayed altered patterns of CDK activity instead of the strong and steady increase in CDK activity in control cells (**Figure 4E**, top rows of “acute” and “proliferative crisis”). After 30 days of chronic cyclin E overproduction, the patterns of cell cycle phases were more typical of control cells, although interestingly, CDK activity was lower in S and G2 phases compared to control cells (**Figure 4E, adaptation, 30-33 days**).

As an additional measure, we examined all the cells in the final frame of each movie by automated image analysis at the end of each 3 days of imaging and plotted the CDK1/2 activity for each cell (**Supplemental Figure S4C**). Based on our prior observations and those of others (Spencer *et al*., 2013; Schwarz *et al*., 2018) cells with a cytoplasmic:nuclear reporter ratio of 0.75 or higher are typically committed to S phase entry or are already in S or G2 phase. We therefore marked the 0.75 ratio as a threshold for high CDK1/2 activity associated with active proliferation. Although activity was high very early (1^st^ 3 hours) after cyclin E induction (**Figure 4D**), by the end of 3 days of cyclin E overproduction, the population with high CDK1/2 activity had already dropped substantially and remained low during the proliferative crisis (**Supplemental Figure S4C, ~ days 9-12**). The fact that we identified some proliferation at all time points argues against the adaptation mechanism being strictly through mutation. We expect that if adapted cells were primarily the descendants of rare mutants, then we would have observed mostly arrested or dead cells and rare clones expanding during and after the proliferative crisis. Instead, we observed dispersed proliferating cells of different proportions throughout the culture at all timepoints.

Taken together, we conclude that chronic cyclin E overproduction induces replication stress and DNA damage that accumulates over many cell divisions and ultimately induces a transient proliferative crisis. During this crisis, individual cells may permanently arrest in either G1 or G2, and cells that arrest in G2 for prolonged periods can skip mitosis and persist in a G1/G0-like state with low CDK activity. A subset of cells continued to proliferate with altered CDK dynamics, and these cells establish a new population that has adapted to cyclin E overproduction.

### Transcriptome shifts in cell cycle-regulated genes during adaptation to cyclin E overproduction

Cells that adapted to cyclin E overproduction proliferated slightly faster than control cells, yet had longer G1 phases and intermediate origin licensing compared to cells during the initial response to cyclin E. We also found that in the absence of cyclin E overproduction, cells maintained intermediate origin licensing through many cell divisions (**Figure 3E**). We hypothesized that adaptation might involve gene regulation changes that allow cells to return to robust proliferation. To identify such changes, we performed RNA sequencing on three independent replicates from key timepoints during adaptation: day 0 (“control” without induction), day 2 (“acute”), day ~9-15 (“proliferative crisis”), or day ~35 (“adapted”) after continuous cyclin E induction. Because adaptation dynamics varied among replicates, we monitored cell proliferation throughout adaptation as in **Figure 3A** and collected RNA from cells when proliferation was slowest and when it had fully rebounded (see Materials and Methods). Additionally, we withdrew doxycycline and allowed cells to recover for another 20 days (“recovery”).

We first examined induction of *CCNE1* mRNA across timepoints and within replicate groups. Although samples were collected independently during adaptations that were initiated weeks apart, the levels of *CCNE1* mRNA were very similar among replicates (**Figure 5A**). After 2 days of acute induction, *CCNE1* mRNA abundance increased more than 20-fold, similar to the 10-fold induction of cyclin E protein by 20 ng/ml doxycycline (e.g. **Figure 1A**, compare lanes 7 and 11). *CCNE1* mRNA levels decreased during proliferative crisis and in adapted cells to an intermediate level that was still significantly higher than endogenous *CCNE1*. The mechanism of this downregulation was not due to loss of the tet repressor mRNA (not shown), but could be epigenetic partial silencing of the transgene, or changes in mRNA processing or stability. We presume this mRNA downregulation after the acute phase is the reason for partial cyclin E protein downregulation in **Figure 3A** (inset, lane 4) and **Figure 4B**. Cells that had reduced expression – despite starting from a clonal population derived from a single transduced cell - may have been selected for during proliferative crisis. As expected, *CCNE1* mRNA returned to endogenous levels after doxycycline withdrawal (**Figure 5A**).

**Figure 5.**
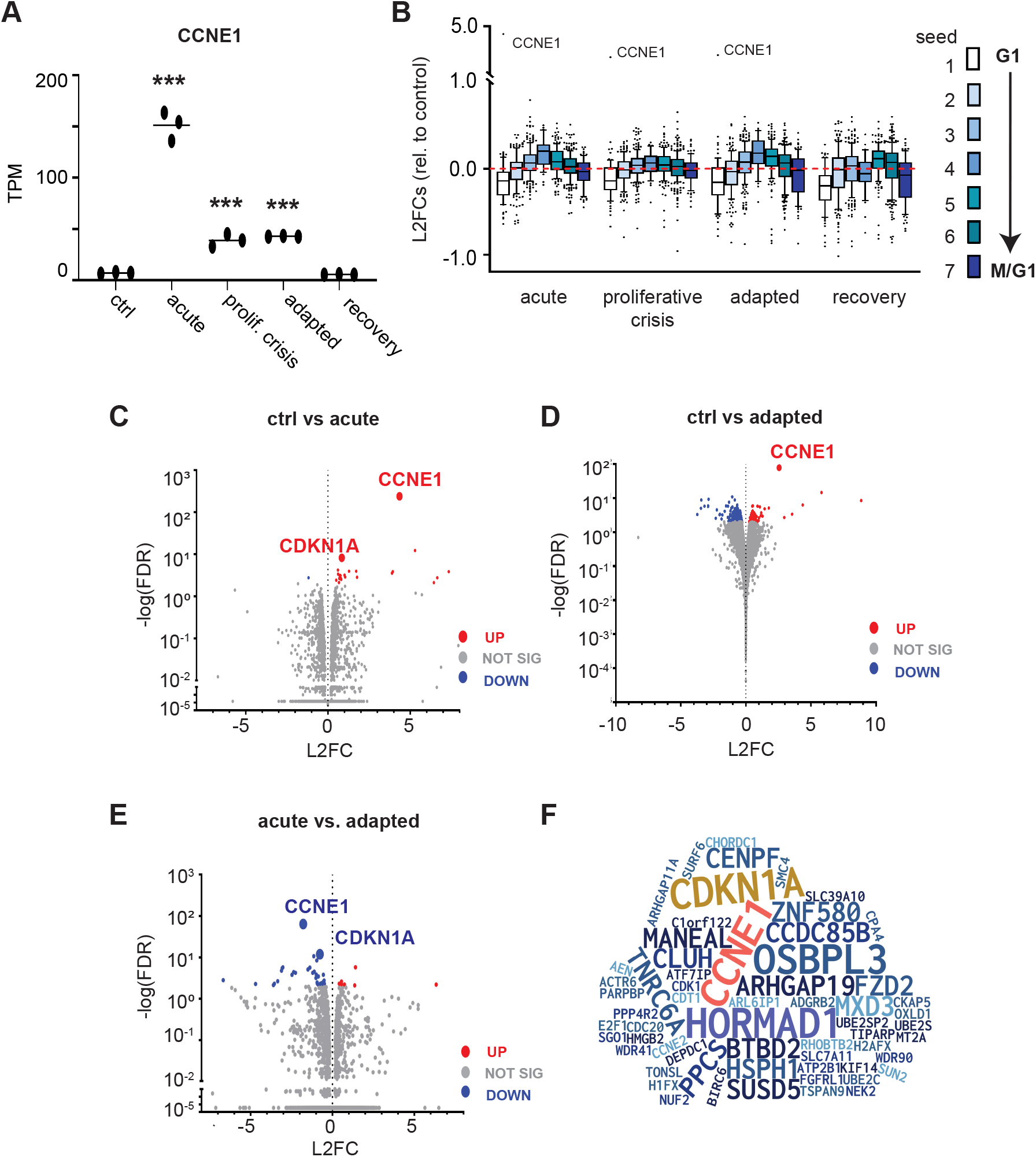
Transcriptome shifts in cell cycle-regulated genes during adaptation to cyclin E overproduction. (A) Cells were harvested for mRNA analysis at the indicated times during cyclin E overproduction (20 ng/ml doxycycline) and two weeks after induction was removed. Ctrl = no dox, acute = 2 days of induction, proliferative crisis = 9-15 days of induction at the point of lowest proliferation, adapted = >30 days of induction, and recovery = 2 weeks after induction was removed from adapted cells. *CCNE1* expression in transcripts per million (TPM) for at each of the 5 timepoints (N=3 independent biological replicates). * indicate adjusted p values of differential expression compared to day 0 for *CCNE1* based on DESEQ. \**p* ≤ 0.05, \*\**p* ≤ 0.005, \*\*\**p* ≤ 0.0005. (B) Log2 fold change (L2FC) relative to uninduced control (day 0) of the genes in seven periodic seed curves defined in Dominguez *et al*. (2016). Seed 1 is genes that peak in G1 and seed 6 and 7 genes peak in M phase and the M/G1 transition; box-plots mark the medians and interquartile ranges. (C) Volcano plot of all genes comparing control to day 2 after induction (acute); statistically significant genes determined by L2FC >2 are marked in red for upregulated genes and blue for downregulated genes. Genes whose change was not statistically significant (L2FC <2) are marked with gray dots. (D) As in C comparing control to adapted cells. (E) As in C & D but comparing day 2 (acute) to day 35 (adapted). (F) Word cloud of all statistically significant genes in any pairwise comparison; letter sizes correspond to frequency of genes in all possible comparisons. See also Supplemental Table 1.

Surprisingly, we detected relatively few significantly differentially expressed genes (DEGs) across time points (we set the cutoff using a -log(false discovery rate) ≥ 2; 426 genes). Additionally, we found high correlations among replicates within each sample (inter-sample Spearman’s r > 0.95, based on all detected mRNAs, see Materials and Methods). Given the robust impact of *CCNE1* induction on cell cycle-associated phenotypes (**Figures 1–3**), we focused our analysis on cohorts of genes whose expression oscillates with the cell cycle that were defined in a prior study (Dominguez *et al*., 2016). We assessed differentially expressed genes relative to time 0 (control cells) for groups of genes assigned to seven periodic seed curves that peak progressively during the cell cycle. Seed 1 genes have peak expression in G1, seed 4 genes peak in G2 phase, and seeds 6 and 7 genes peak in M phase and the M/G1 transition respectively. We then plotted the log2 fold changes (inductions vs. non-induced) for each of the seven gene cohorts at each time point. From these analyses, we found that the G1 cohort (seed 1, white boxes) was clearly downregulated in cyclin E-overproducing cells with a corresponding increase in seed 4 (G2) genes (**Figure 5B**). Expression of seed 1 genes remained low even after doxycycline withdrawal. These shifts in expression, particularly for G1-expressed genes, may reflect cell cycle phase length changes, may be the driving cause of those changes, or both.

The *CDKN1A* gene encoding the CDK inhibitor p21 was among the few genes whose expression changed significantly in comparisons between control and day 2 (acute overproduction, **Figure 5C**), and between day 2 and day 35 (acute vs. adapted cells, **Figure 5E**). Elevated *CDKN1A* mRNA is consistent with the elevated 53BP1 nuclear bodies and p21 protein levels during both the acute response and proliferative crisis when markers of replication stress were highest (**Figure 4A and 4B**). Interestingly, *HORMAD1* (which encodes cancer/testes antigen gene 46) was also found to be significantly up-regulated in our sample set. *HORMAD1* is involved in multiple DNA-dependent processes during meiosis, its expression is normally restricted to germ cells (Simpson *et al*., 2005).Even moderate *HORMAD1* misexpression in these somatic epithelial cells represents a large fold-increase. Despite this increase, the absolute levels of *HORMAD1* mRNA were still quite low, and we were unable to detect HORMAD1 protein in adapted cells (not shown).

To investigate potential operant pathways involved in adapting to cyclin E overproduction, we performed Gene Set Enrichment Analysis (GSEA) with our transcriptome data. Acute cyclin E overproduction caused downregulation of several signaling pathways, and the estrogen response late and early pathways were predominant as down-regulated pathways during proliferative crisis and adaptation (**Supplemental Figure S5A**). We found few up-regulated pathways to be statistically significant across time point comparisons (**Supplemental Figure S5B**). In addition, few statistically significant individual genes were upregulated between control and proliferative crisis cells (**Supplemental Figure S5C**). However, when we compared all genes at the beginning (control) and end of the adaptation, we found that selected genes involved in origin licensing (*MCM5, CDT1*) were down-regulated and DNA damage genes were either up-regulated (*ATM*) or down-regulated (*H2AFX*), though these individual gene changes fell short of our criteria for statistical significance (**Supplemental Figure 5D**). Lastly, to better visualize the impact on cell cycle genes over the course of adaptation we constructed a word cloud of significantly changed cell cycle genes weighted on the frequency of their changes in all possible pairwise comparisons (**Figure 5F**); all significant gene expression changes relative to controls are provided in **Supplemental Table 1**.

### Parallel gene expression changes in breast cancers with high *CCNE1* expression

We next assessed *CCNE1* expression in human breast tumors (The Cancer Genome Atlas, TCGA). As expected, *CCNE1* mRNA correlates with *CCNE1* copy number changes, and tumors harboring *CCNE1* amplifications expressed ~20-fold higher *CCNE1* mRNA than diploid tumors (**Figure 6A**, Materials and Methods). This level of *CCNE1* overexpression in primary tumor samples is similar to the fold-induction of *CCNE1* achieved in our inducible RPE1-hTert model. Moreover, markers associated with the DNA damage response, such as phosphorylated CHEK1, also correlated with *CCNE1* expression in the TCGA database (**Figure 6B**) and in another analysis of human biopsies (Guerrero Llobet *et al*., 2020), consistent with elevated cyclin E-induced replication stress. We are thus confident that we had recapitulated at least some of the consequences of cyclin E overproduction that are visible in human tumors. This experimental approach to analyze the process of adaptation to one oncogene can shed light on both the early stress induced by oncogene expression and the changes that arise to accommodate that stress.

**Figure 6.**
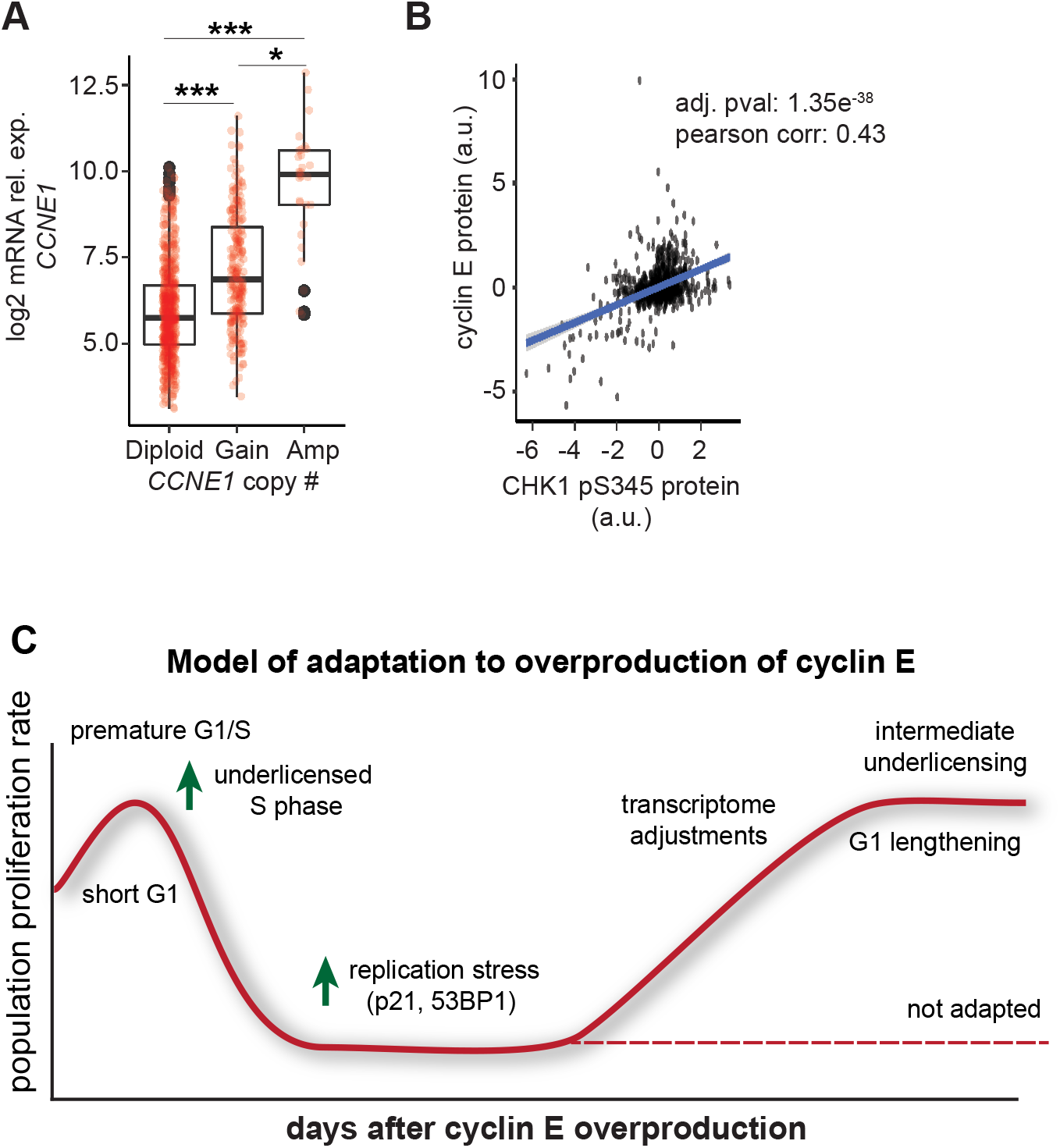
Cyclin E overproduction in RPE1-hTERT cells is representative of cyclin E-overexpressing breast cancers. (A) *CCNE1* expression relative to gene copy change in TCGA breast tumors. Number of cases in TCGA for *CCNE1:* Diploid (601), Gain (232); Amplification (29) \**p* ≤ 0.05, \*\*\**p* ≤ 0.0005. (B) Phospho-CHEK1 relative to cyclin E protein in human breast tumors from the TCGA database. (C) Model for adaptation to cyclin E overproduction: The initial response is a short G1 which leaves insufficient time for origin licensing. These cells experience replication stress, accumulate DNA damage, and slow or arrest proliferation. With continued cyclin E overproduction, a subset of cells eventually adapt, lengthen G1 phase, and proliferate with perturbed cell cycle gene expression.

## DISCUSSION

Cyclin genes are frequently overexpressed in human cancers either from copy number changes or loss of normal upstream signaling control (Donnellan and Chetty, 1999; Chu *et al*., 2021). The effects of cyclin overproduction on proliferation and genome stability are not easy to dissect in fully developed cancer cells however, because the cyclin changes are combined with many other genetic and epigenetic changes that activate growth signaling, inhibit apoptosis, and impair checkpoints (Hanahan and Weinberg, 2011). We analyzed cells throughout the process of adaptation for changes in cell cycle phase length, markers of replication stress, and CDK activity using a variety of methods at single-cell resolution. This approach differs from comparing starting cell populations with resistant or fully-adapted populations.

To isolate the effects of G1-S cyclin overproduction from these other changes, we produced each cyclin under inducible control in non-transformed epithelial cells. Cyclin D1 overproduction induced slightly faster proliferation and shorter G1 phases but did not cause a proliferative crisis in the same way that cyclin E overproduction did. Why is cyclin E particularly toxic? Cyclin E induced a much shorter G1 phase than either cyclin D or cyclin A did. There may be a threshold of G1 length below which cells cannot proliferate well. Cyclin E overproduction triggered premature S phase entry, in part, because unlike cyclin D, direct cyclin E/CDK2 substrates include those required for origin firing (Boos *et al*., 2011; Kumagai *et al*., 2011; Siddiqui *et al*., 2013). The effects of cyclin D overproduction may mimic growth factor signaling pathways but only indirectly induce S phase entry. Cyclin A does not appear to be rate-limiting for S phase entry in these cells/under these conditions, perhaps because it’s difficult to increase cyclin A levels during G1 because of degradation by APC/C (Geley *et al*., 2001). The capacity of cyclin A to shorten G1 phase in these cells, therefore, appears more limited.

The strong proliferative crisis induced by cyclin E overproduction is uniquely associated with substantial deficiencies in origin licensing at the time of premature S phase entry. We propose that a major contributor to the replication stress in cyclin E-overproducing cells is insufficient origin licensing. We acknowledge that other mechanisms may also contribute to cyclin E-induced replication stress. For example, the premature S phase can also induce replication-transcription collisions by allowing intragenic origins to fire that normally would not (Macheret and Halazonetis, 2018). Previous studies have also noted less MCM loading in cyclin E-overproducing cells, but these results didn’t examine long-term proliferation effects and have been interpreted within the paradigm of direct CDK-mediated inhibition of MCM loading (Ekholm-Reed *et al*., 2004). The direct inhibition paradigm was first established in studies of budding and fission yeasts where MCM loading proteins are inhibited by CDK-mediated phosphorylation (Nguyen *et al*., 2001; Diffley, 2004). In mammalian cells MCM loading proteins are also inhibited during S phase to prevent inappropriate re-licensing, but interestingly, the mechanisms of that inhibition may be more from cyclin A-mediated phosphorylation than from cyclin E, at least at normal cyclin expression levels. For example, CDT1 is degraded by the SCF^SKP2^ E3 ubiquitin ligase after phosphorylation, but CDT1 only binds cyclin A and not cyclin E (Zhou *et al*., 2020); cyclin E is also not required for Cdt1 degradation during replication in *Xenopus laevis* extracts (Arias and Walter, 2005). Similarly, the largest subunit of ORC, Orc1, is degraded in S phase but is a much better substrate for cyclin A/CDK2 than it is for cyclin E/CDK2 (Méndez *et al*., 2002). Human CDC6 is translocated to the cytoplasm during S phase after phosphorylation mediated by cyclin A, but not by phosphorylation at serine 54 which is attributed to cyclin E (Petersen *et al*., 1999; Yim *et al*., 2013). Importantly, cyclin E activates rather than inhibits mammalian CDC6 in the sense that cyclin E-dependent phosphorylation at serine 54 stabilizes CDC6 during late G1 phase (Mailand and Diffley, 2005). We assert, that unlike yeast CDKs that directly inhibit MCM loading proteins, mammalian cyclin E is not a strong direct inhibitor of origin licensing, and that MCM loading factor inactivation is largely dependent on other CDKs.

Cyclin E/CDK2 activity rises in late G1 phase at the time that origin licensing is most active (Mailand and Diffley, 2005), and so it would be counterproductive for cyclin E to block origin licensing in G1. Cyclin A on the other hand is expressed only once S phase has started. The strong underlicensing we demonstrate in cyclin E-overproducing cells is more likely attributable to the *indirect* effects of premature S phase onset. Activating DNA replication triggers CDT1 degradation once PCNA is loaded at replication forks and recruits the CRL4^CDT2^ E3 ubiquitin ligase (Arias and Walter, 2006; Hu and Xiong, 2006; Senga *et al*., 2006). Since CDT1 is essential for MCM loading (Maiorano *et al*., 2000), early S phase onset from cyclin E overproduction stops all further MCM loading. This degree of underlicensing then causes accumulating replication stress that induces the proliferative crisis we document here. Of note, cyclin A overproduction had minimal effects on G1 length (**Figure 1C**) and induced extremely modest underlicensing (**Figure 2D and Supplemental Figure 1F**). This limited licensing inhibition could have resulted from direct MCM loading factor inhibition in G1 by cyclin A-mediated phosphorylation. We also consider the possibility that at high levels, cyclin E may inactivate MCM loading proteins directly which could contribute to the moderate underlicensing in adapted cells. One prior study identified a CDK2 phosphorylation site in MCM7 that may interfere with MCM2-7 complex formation and chromatin loading, and that site is phosphorylated by both cyclin E and cyclin A *in vitro* (Wei *et al*., 2013). Our findings add nuance to a fundamental understanding of the relationship between CDK activity and licensing in mammals and the importance of G1 length to avoid replication stress.

How then did cells overcome the cyclin E-induced replication stress and return from crisis to robust proliferation? Remarkably, adaptation to high ectopic cyclin E was not simply a reversal of the overproduction itself. Adapted cells still overproduced cyclin E, and cyclin E-associated kinase activity stayed elevated relative to the starting population. If a major cause of the replication stress was underlicensing due to truncating G1 phase, then the longer G1 phases that developed in adapted cells likely provided enough time for adequate origin licensing. Indeed, adapted cells were substantially less underlicensed than cells experiencing acute cyclin E-overproduction. These observations are generally consistent with high cyclin E-expressing cancer cell lines proliferating well, although even higher levels of ectopic cyclin E suppressed licensing in the first G1 after induction (Ekholm-Reed *et al*., 2004; Asghar *et al*., 2017; Geng *et al*., 2018).

The mechanism(s) by which G1 lengthened may have included generally down-regulating the cohort of genes whose expression normally peaks in G1 phase. How these G1 genes were down-regulated is not yet clear, but it’s possible that one or more components of the upstream signaling and regulatory systems that induce these genes are less active; the estrogen response pathway was significantly down-regulated in adapted cells. The adjustments in adapted cells appear to have occurred to many genes by a small amount each rather than by dramatic changes in a handful of genes. Thus far, we have analyzed transcriptome changes, but future studies of proteome changes will shed additional light on these mechanism(s) of adaptation. Of additional relevance, not all cells with short G1 phases are underlicensed. A hallmark of pluripotency is a very short G1 phase, but we previously demonstrated that pluripotent stem cells license origins very fast and achieve high levels of MCM loading by the onset of S phase (Matson *et al*., 2017). Our adapted populations did have improved origin licensing compared to cells with acute cyclin E overproduction, but also longer G1 phases, not typically associated with faster origin licensing.

In addition to reduced G1-regulated gene expression, the induction of DNA damage and replication stress markers (53BP1 and p21) almost certainly contributed to the proliferative crisis, and their down-regulation contributed to eventual adaptation. It is not surprising that p21 levels were induced (an 8-fold increase between control vs. day 12 cells) in cells experiencing high replication stress and proliferative crisis since it is a well-established component of the DNA damage response (El-Deiry *et al*., 1993; Macleod *et al*., 1995). The overall increased p21 protein levels in fully adapted cells (**Figure 4B**) may also have contributed to G1 lengthening. Although overall cyclin E-associated kinase activity remained elevated compared to controls as measured via immunoprecipitation (**Figure 3C**), the dynamics of CDK activation in individual cells was clearly perturbed and appeared more delayed (**Figure 4E**).

Adaptation in the evolutionary sense is typically thought of as a selection for genetic changes. The adaptation we characterized here is unlikely to be driven by mutations, although mutations certainly may have occurred. Indeed, a previous study of transient cyclin E overproduction documented genomic deletions in individual cells (Teixeira *et al*., 2015b). On the other hand, cells chronically overproducing cyclin E in our study did not appear to adapt by the expansion of rare mutant clones, but rather by non-genetic perturbations. We cite three observations in support of this interpretation. First, we routinely inspected cultures during proliferative crisis and return to rapid proliferation, and we readily found proliferating cells at all time points rather than widespread arrest combined with rare proliferating clones. Second, cultures consistently overcame proliferative crisis within two to three weeks. We derived adapted populations starting from both large and small cell numbers (as few as 20,000 cells). A mutation mechanism would presumably be sensitive to the size of the starting population and rare enough to affect the timing of recovery from crisis. Third, adapted cells had intermediate origin underlicensing, and returning cyclin E levels to normal resulted in a gradual reversion to full licensing over a period of weeks. A mutation would not revert in the absence of any evidence for selective pressure. We acknowledge that contributions from mutations cannot be completely ruled out, particularly as cells return to proliferation from crisis. However, adaptation in the evolutionary sense is also thought to be slow and gradual. Our system yielded phenotype changes over a period of weeks. Instead, we speculate that the primary mechanism is epigenetic, signaling rewiring, post-transcriptional/translational adjustments or some mosaic combination of these or other non-genetic changes.

In conclusion, we report here a quantitative and thorough characterization of epithelial cell adaptation to replication stress caused by cyclin E overproduction. This study is unique in that it characterizes adaptation throughout the process at single-cell resolution in contrast to comparing the starting and adapted cell populations. Adaptation was surprisingly and consistently rapid and resulted in cells with altered growth and replication phenotypes. We propose a model for adaptation to cyclin E overproduction (**Figure 6C**). First, cells experience high CDK activity which causes a short G1 and underlicensing. The underlicensing results in accumulating replication stress coupled with DNA damage which induces a period of slowed proliferation with cell cycle patterns and arrests that vary among individual cells. Over the course of several more cell generations, the proliferating population has longer G1 phases, shifts in gene expression, intermediate licensing, and less DNA damage. This study visualizes the plasticity of the cell cycle while providing a careful and quantitative examination of the impact of cyclin E on the cell cycle over many cell generations. In addition, the experimental approach described here may recapitulate events during oncogenesis as cells accommodate replication stress from oncogene activation.

## MATERIALS AND METHODS

### Cell culture & cell line construction

RPE1-hTERT cells were verified by STR profiling (ATCC, Manassas, VA) and confirmed to be mycoplasma negative. Cells were maintained in Dulbecco’s modified Eagle medium (DMEM) supplemented with 10% fetal bovine serum (FBS), 2 mM L-glutamine, 1x pen/strep & incubated at 37°C in 5% CO_2_. Cells were passaged with 1x trypsin prior to reaching confluency. RPE1-hTERT cells with the pINDUCER20-cyclin constructs were passaged every 3 days. To package lentivirus, pINDUCER20 (Addgene plasmid #44012) with human cyclin E1, cyclin D1, or cyclin A2 cDNAs were co-transfected with ΔNRF and VSVG plasmids (gifts from Dr. J. Bear, UNC) into HEK293T cells with 50 μg/mL polyethylenimine (PEI)-Max (Aldrich Chemistry). RPE1-hTERT cells were transduced with the viral supernatant from the transfections in the presence of 8 ug/mL polybrene (Millipore, Burlington, MA) for 24 hr, then selected with 500 ug/mL geneticin (Gibco # 10131035). Clones from the polyclonal population were isolated and screened for inducible expression of the respective protein (cyclin D1, cyclin E1, or cyclin A2).

Construction of reporter cell lines: PEI-Max was used to transfect PCNA-mTurquoise (Grant *et al*., 2018), DHB-mCherry (gift from S. Spencer, University of Colorado-Boulder, Boulder, CO (Spencer *et al*., 2013)) into 293T cells using previous methods as stated above to package virus. The virus generated was then used to transduce RPE1-hTert cells already containing the pINDUCER20-cyclin E1 construct using the manufacturers recommended protocol. Cells with stable integration of the plasmids were selected using 500 μg/mL G418 (Gibco). Colonies from single cells were selected by visual inspection for even expression of fluorescently tagged proteins.

### Cloning

cDNAs were subcloned into the pINDUCER20 construct (Meerbrey *et al*., 2011) using either the Gateway cloning method (Life Technologies) or Gibson Assembly following protocols previously described (Matson *et al*., 2017; Grant *et al*., 2018). PCR fragments were amplified with Q5 polymerase (New England Biolabs, NEB, # M0491L). DNA fragments were isolated using the Qiaprep spin miniprep kit (Qiagen). Plasmids were transformed into DH5α *Escherichia coli* strains. pENTR constructs were then combined with pINDUCER20 (Addgene plasmids#44012). Plasmids were validated via sequencing (Eton Biosciences) for the desired insert using appropriate primers.

### Immunoblotting

Cells were lysed with CSK buffer (300 mM sucrose, 300 mM NaCl, 3 mM MgCl^2^, 10 mM PIPES pH 7.0) containing 0.5% Triton X-100 supplemented with protease and phosphatase inhibitors (0.1 mM AEBSF, 1 μg/ mL pepstatin A, 1 μg/ mL leupeptin, 1 μg/ mL aprotinin, 10 μg/ ml phosvitin, 1 mM β-glycerol phosphate, 1 mM Na-orthovanadate) for 20 min on ice. Lysates were centrifuged at 4°C for 10 min at maximum speed in a microcentrifuge. Supernatants were removed and lysates were diluted with SDS loading buffer (final: 1% SDS, 2.5% 2-mercaptoethanol, 0.1% bromophenol blue, 50 mM Tris pH 6.8, 10% glycerol) and boiled. Samples were separated on appropriate SDS-PAGE gels, and proteins transferred onto nitrocellulose (GE Healthcare, Chicago, IL) or polyvinylidene difluoride (PVDF) membranes (Thermo Fisher, Waltham, MA). Following transfer, total protein was detected by staining with Ponceau S (Sigma-Aldrich). Membranes were blocked for 1 hour at room temperature in 5% milk or 5% BSA in Tris-Buffered-Saline-0.1% Tween-20 (TBST). Membranes were then incubated in primary antibody for 16-18 hours at 4°C with constant shaking in either 2.5% milk or 5% BSA in TBST with 0.01% sodium azide. After washing in TBST, membranes were incubated with horseradish peroxidase-conjugated secondary antibody in either 2.5% milk or 5% BSA in TBST for 1 hour at room temperature and washed with TBST. Signals were detected using ECL Prime (Amersham, United Kingdom) and a ChemiDoc MP (Bio-Rad). This imaging system indicates any signals above the linear range of detection, and we did not include any images with signals outside that range. Antibodies used for immunoblotting were: cyclin E1 (Cell Signaling Technologies, Beverly, MA, cat# 4129), cyclin D1 (Santa Cruz Biotechnology, cat# sc753), cyclin A2 (Cell Signaling Technologies, Beverly, MA, Cat# 4656). Secondary antibodies used included: anti-rabbit IgG HRP-conjugated (1:10,000, Jackson Immuno Research) & goat anti-mouse IgG HRP-conjugated (1:10,000, Jackson Immuno Research).

### Immunoprecipitation

Cells were collected and frozen, then resuspended in Kischkel buffer (50 mM Tris pH 8.0, 150 mM NaCl, 5 mM EDTA, 1% Triton X-100) supplemented with 100 nM ATP and protease/phosphatase inhibitors as used for immunoblotting above for 20 min on ice. Cells were centrifuged at 4 °C for 10 min. at 15,000 rpm and the supernatants removed. Lysates were pre-cleared for 45 min with magnetic beads (Invitrogen, Dynabeads protein G, cat# 10003D) and nuclear digestion buffer (10 mM HEPES pH 7.9, 10 mM KCl, 1.5 mM MgCl2, 340 mM sucrose, 0.1 mM glycerol) + 1 mM ATP + protease/phosphatase inhibitors (used above). Following quantification via Qubit or Bradford assay (Bradford, 1975), precleared lysates were adjusted to 1X high stringency IP buffer (Active Motif, cat# 37510) with protease/phosphatase inhibitors, 0.01 mM ATP, & detergent (Active Motif, cat# 37517). Lysates were then incubated with beads that had been pre-incubated with antibody in cold blocking buffer (1X PBS + protease/phosphatase inhibitors + ATP) for 6-8 hours rotating at 4 °C, then washed to remove unbound antibody. Lysates and antibody-bound beads were incubated with rotation at 4 °C for 16-18 hour. Unbound protein was removed the next day, beads were washed 3x with 5x high stringency IP buffer (diluted to 1X) supplemented with protease/phosphatase inhibitors + ATP and split 50% for immunoblotting and 50% for kinase assay. Antibodies used: cyclin E1 (Invitrogen, cat# 32-1500) & cyclin E (Cell Signaling Technologies, cat# 4129).

### Protein kinase assay

Protein-antibody-bead combinations were resuspended in kinase buffer (50 mM Tris pH 7.5, 10 mM MgCl2, and 0.01 mM ATP, + protease/phosphatase inhibitors). 2 μCi of [γ-^32^P]ATP was added per sample, + 1 μg histone H1 protein (Abcam, cat# ab198676). Samples were incubated for 30 min. at 30 °C. 20% 2-mercaptoethanol + sample buffer was added to each sample, boiled for 5 min, centrifuged 15,000 rpm for 30 sec, then separated by SDS-PAGE, and the gel was stained with Coomassie blue, dried and exposed to a PhosphorImaging cassette (GE Healthcare). The phosphorylated histone H1 signal was quantified by subtracting the serum control from all experimental signals, then all signals within one experiment were normalized to 1 based on activity in the control (endogenous cyclin E) sample.

### Flow cytometry

For analytical flow cytometry, EdU (Santa Cruz Biotechnology) was added to 10 μM 30 min prior to collection with trypsin. Cells were permeabilized with 500 μL cytoskeletal (CSK) buffer supplemented with 0.5% Triton X-100 and protease and phosphatase inhibitors as used above for immunoblotting, then fixed with paraformaldehyde and subjected to antibody staining and EdU detection as described here. http://dx.doi.org/10.17504/protocols.io.bba8iihw. Samples were analyzed on an Attune NxT flow cytometer (Life Technologies). Flow cytometry data were evaluated using FCS Express 7.0 software (De Novo, Glendale, CA). Cells were gated on FS(A) x SS(A) plots, and singlets were gated using DAPI(A) x DAPI(H) plots. Control samples were used as described previously (Matson *et al*., 2017). This sample was treated the same as other samples by incubating with azide (either 647 or 488) and the appropriate secondary antibody. The following antibody/fluorophore combinations were used: (1) MCM (measuring origin licensing): Alexa 647-azide (Life Technologies), primary: Mcm2 (BD Biosciences, cat# 610700), secondary: donkey anti-mouse-488 (Jackson Immuno Research), DAPI. (2) primary: γ-H2AX (measuring DNA damage): Alexa 488-azide (Life Technologies), primary: γ-H2AX (Cell Signaling Technologies, Beverly, MA, cat# 9718); secondary: donkey anti-rabbit 647 (Jackson ImmunoResearch), DAPI (Life Technologies, cat# D1306).

### Doubling time

Doubling times were calculated by plating cells and counting cell numbers using a Luna II automated cell counter (Logos Biosystems, South Korea) 72 hours after plating. Each timecourse was repeated a minimum of 3 times. P = final # of cells, P0 = initial # of cells PD = population doubling calculated as log2(P/P0), and Dt = population doubling time is calculated as T (time) / PD. A minimum of 3 biological replicates were averaged for a final mean doubling time and graphed as 1/Dt. GraphPad Prism and Excel were used to plot final graphs.

### Live cell imaging

RPE1-hTERT cells were plated as single cells in 24-well glass-bottom dishes (Cellvis) at a density of 20,000 cells/well. Cells were grown in FluoroBrite media (Gibco, #A18967-01) with 10% fetal bovine serum (FBS), 2 mM L-glutamine, 1x pen/strep at 37°C in a humidified enclosure with 5% CO_2_ (Okolab) and stimulated with 20 ng/mL doxycycline where indicated. Images were collected starting 4 hours after plating, and cells were imaged for 72 hours with images collected every 10 minutes using a Nikon Ti Eclipse inverted microscope (20x 0.75 NA dry objective lens) with the Nikon Perfect Focus system. Images were captured using an Andor Zyla 4.2 sCMOS detector with 12-bit resolution. All filter sets were from Chroma, CFP - 436/20 nm; 455 nm; 480/40 nm (excitation; beam splitter; emission filter), YFP - 500/20 nm; 515 nm; 535/30 nm; and mCherry - 560/40 nm; 585 nm; 630/75 nm. Images were collected with the Nikon NIS-Elements AR software.

### Cell tracking

Cells were tracked using ImageJ software (https://imagej.nih.gov). Custom Python scripts (v3.7.1) in Jupyter Notebooks (v6.1.4) were used following tracking to perform single cell biosensor analyses. Individual cells were tracked and segmented in movies (time-lapse experiments) using a previously described user-assisted approach (Grant *et al*., 2018).

### Immunofluorescence microscopy

Cells were fixed with 4% paraformaldehyde (PFA) for 15 min., permeabilized with 0.5% Triton X-100 for 30 min., then incubated with primary antibody (anti-53BP1, Novus Biologics, #NB100-304) at 4°C. The next day cells were incubated with secondary anti-rabbit antibody conjugated to Alexa Fluor-488 for 1 hour at room temperature. Following washing, cells were incubated with 1 μg/mL DAPI and imaged with appropriate filters. Slides were scanned with a Nikon Ti Eclipse inverted microscope (details in Live cell imaging section) with a 40x 0.95 NA objective. For each time point 20 fields of view were manually selected for analysis yielding between 250 and 1500 cells to be analyzed. Nuclei were detected based on DAPI signal using Python implementation of the Stardist segmentation algorithm (Weigert *et al*., 2020). 53BP1 nuclear bodies were detected using Difference of Gaussians image enhancement algorithm followed by a watershed segmentation algorithm using Scikit-image (van der Walt *et al*., 2014) in Python.

### Statistical analysis

Bar graphs represent means, and error bars indicate standard error of the mean (SEM), unless otherwise noted. The number and type of replicates are indicated in the figure legends. Significance tests were performed using a one-way ANOVA test, as indicated in the figure legends, unless otherwise specified. Statistical significance is indicated as asterisks in figures: * p ≤ 0.05, ** p ≤ 0.05, *** p ≤ 0.005 and **** p ≤ 0.0005. GraphPad Prism v.8.0 and Python were used for statistical analysis.

### RNA extraction, library preparation and RNA sequencing

RNA was extracted from RPE1-hTERT cells using the Quick-RNA Miniprep Kit (Zymo Research, Irvine, CA, USA) with chloroform extraction. Libraries for each sample were prepared, at the same time, using KAPA Hyperprep mRNA Library Kit (Roche) and Illumina adapters (New England Biolabs) starting with 1 μg RNA for each sample. Pooled libraries were sequenced with a Nextseq 500 high output 75 cycle kit for single-end reads.

To analyze RNA sequencing data, Kallisto (v0.44.0) was used to quantify transcript abundance against a reference human genome (hg38) followed by generation of transcripts per million (TPM) tables. Kallisto output was imported into DESeq2 (v1.24.0) to determine differential gene expression between time points. Ranked lists of genes with differential expression between timepoints based on log2 fold change were used for gene set enrichment analysis (GSEA). Pairwise correlations were performed using detected genes with TPM of at least 5. Cell cycle gene sets were derived from RNA sequencing (Dominguez *et al*., 2016) and in total 931 genes overlapped between with this dataset. Periodic “seeds” 1-7 from (Dominguez *et al*., 2016) were used to investigate expression changes in this study. Breast cancer (BRCA) data from Tumor Cancer Genome Atlas Data were downloaded from the Cancer Bioportal Repository (Gao *et al*., 2013). Copy number variation and gene expression values (RSEM+1) were used from the published datasets. Statistical significance tests are denoted in figure legends with p value adjustment when multiple comparisons were made.

## Supporting information

Limas supplemental figures 1-6

Limas Supplemental Table 1

## DATA AVAILABILITY

The data discussed in the paper are included in the figures and supplements. RNA-Seq data has been deposited in the Gene Expression Omnibus (GEO) database (accession no. GSE171845).

## SUPPLEMENTARY DATA

Supplementary Figures 1-6 and Supplementary Table 1 are provided.

## ACKNOWLEDGEMENTS

We thank Sidra Qayyum, Megan Justice, Cyrus Vaziri, Michael Emanuele, Lee Graves, & Wayne Stallaert for technical assistance. We thank Jeffrey Jones for lab management. We thank Samuel Wolff and Jeremy E. Purvis for expertise in live cell fluorescence microscopy. The results shown here are in whole or part based upon data generated by the TCGA Research Network: https://www.cancer.gov/tcga.

## Author contributions

Conceptualization: J.C.L., J.G.C., K.M.K.; Validation: J.C.L., K.M.K., M.M.A., D.D.; Formal analysis: J.C.L.,, K.M.K., M.M.A., D.D., A.W.; Investigation: J.C.L., A.N.L., K.M.K., J.G.C., A.M.H, D.F., B.L.M., B.M., M.M.A., D.D.; Writing (first draft): J.C.L., J.G.C., A.N.L., K.M.K., A.W., D.D., M.M.A.; Writing (editing): J.C.L., J.G.C., K.M.K., M.M.A., D.D.; Supervision: J.C.L., J.G.C., Funding acquisition: J.G,C., K.M.K.; Project administration: J.G.C., J.C.L.

---

**Supplemental Table 1. Statistically significant up- and down-regulated genes**.

All genes whose changes were statistically significant in pairwise comparisons. Each tab lists gene names, log2 fold change (L2FC), the -log of the false-discovery rate (FDR), and the direction of the change.

**Supplemental Figure 1. Modest underlicensing induced by cyclin D or cyclin A overproduction**. (A) Gating strategy and color coding for cell cycle phases, MCM loading, and MCM loading just in early S phase on EdU/DAPI plots. (B) Defining gates for MCM antibody negative (“AB-“) cells (gray), G1 cells (blue), mid-to-late S & early S phase cells (orange and green, respectively), and G2-M cells (gray). (C) Licensing in cells after inducing cyclin D expression with 100 ng/ml doxycycline for 48 hours; brackets indicate early S phase cells. (D) Bound MCM intensity in early S phase cells from (C) plotted as histograms (left) and single cells (right). (E) and (F) As in (D) and (C) except cells were induced to overproduce cyclin A with 100 ng/ml doxycycline for 48 hours. Data shown are representative of a minimum of three biological replicates. Histograms in D and F are identical to those in Figure 2D.

**Supplemental Figure 2. Replication stress by Ɣ-H2AX staining**. Gating strategy for measuring replication stress: antibody negative (AB-) cells (gray) and Ɣ-H2AX positive (red). Left: control cells. Right: cells treated 48 hours with 1 nM gemcitabine (gem).

**Supplemental Figure 3. Adaptation to chronic cyclin D or cyclin A overproduction**. (A) Proliferation rate for uninduced control cells (black) and cells overproducing cyclin D1 (100 ng/ml, green) or cyclin A2 (100 ng/ml, blue) and cyclin E1 (20 ng/ml, red, from Figure 3A). (B) Immunoblot analysis of cyclin D1 (top) and cyclin A2 (bottom) after 2 or 39 days of proliferation under constant inducing conditions, lanes 2 and 4. Lane 3 is cells after chronic cyclin D1 or cyclin A2 overproduction 2 days after dox removal.

**Supplemental Figure 4. DNA damage markers and CDK activity during adaptation to cyclin E overproduction**. (A) Representative immunofluorescence images of cells fixed at three-day intervals during adaptation to cyclin E overproduction. DNA damage was scored as bright 53BP1 nuclear bodies (green), and DNA was stained with DAPI (blue). (B) Cells were induced with 20 ng/mL doxycycline to overproduce cyclin E for 2 days or after adaptation (30+ days), then treated with the indicated concentrations of NCS for 48 hours. Endogenous p53, p21, and vinculin (as a loading control) in whole cell lysates were detected by immunoblotting. (C) CDK reporter activity at the end of each 72 hours of live imaging during the course of adaptation. The line at 0.75 marks a threshold of CDK reporter activity above which cells are committed to S phase, in S, or G2 phase.

**Supplemental Figure 5. Individual gene and pathway changes during adaptation to cyclin E overproduction**. (A) Heat map of down-regulated pathways in the indicated pairwise comparisons using GSEA shaded by p-value. Statistically significant pathway changes defined by nominal p-value score <0.05 are highlighted in black with an “S” (light yellow boxes indicate no data) (B) Heat map of up-regulated pathways in the indicated pairwise comparisons using GSEA shaded by p-value. Statistically significant pathway changes are highlighted in black with an “S” (light yellow boxes indicate no data). (C) Volcano plot of all detected mRNAs comparing control to proliferative crisis (day 0 vs ~days 9-12) after induction; statistically significant upregulated or downregulated genes are marked in red or blue respectively. Gene expression changes below the threshold for significance are marked with gray dots. (D) As in C comparing control to recovered cells after doxycycline withdrawal; selected origin licensing and DNA damage response genes are highlighted. See also Supplemental Table 1.

**Supplemental Figure 6. Alternative exposures from Figures 1A, 1D, and 3B**. (A) Lighter exposures from Figure 1A (B) Lighter exposure of Figure 1D (C) Lighter exposure of Figure 3B.

