## Supplementary material for "Quantitative Profiling of Adaptation to Cyclin E Overproduction": Limas supplemental figures 1-6

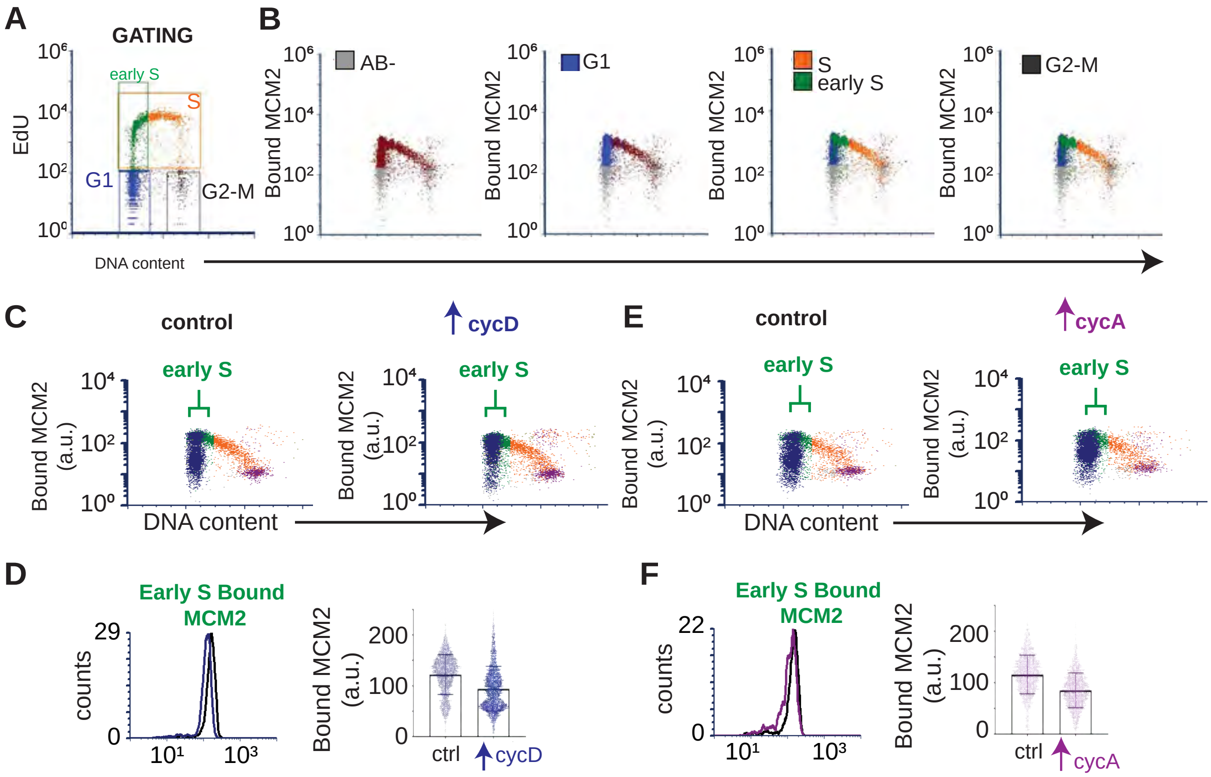


**Supplemental Figure 1. Modest underlicensing induced by cyclin D or cyclin A overproduction**. (A) Gating strategy and color coding for cell cycle phases, MCM loading, and MCM loading just in early S phase on EdU/DAPI plots. (B) Defining gates for MCM antibody negative ("AB-") cells (gray), G1 cells (blue), mid-to-late S & early S phase cells (orange and green, respectively), and G2-M cells (gray). (C) Licensing in cells after inducing cyclin D expression with 100 ng/ml doxycycline for 48 hours; brackets indicate early S phase cells. (D) Bound MCM intensity in early S phase cells from (C) plotted as histograms (left) and single cells (right). (E) and (F) As in (D) and (C) except cells were induced to overproduce cyclin A with 100 ng/ml doxycycline for 48 hours. Data shown are representative of a minimum of three biological replicates.

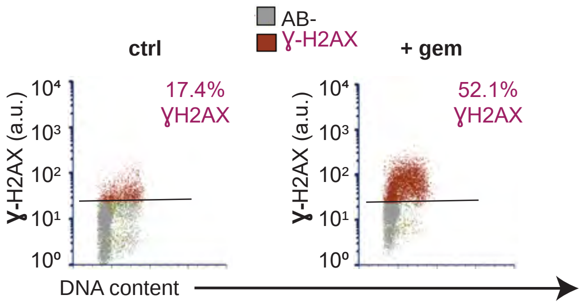


**Supplemental Figure 2. Replication stress by ɣ-H2AX staining.** Gating strategy for measuring replication stress: antibody negative (AB-) cells (gray) and ɣ-H2AX positive (red). Left: control cells. Right: cells treated 48 hours with 1 nM gemcitabine (gem).

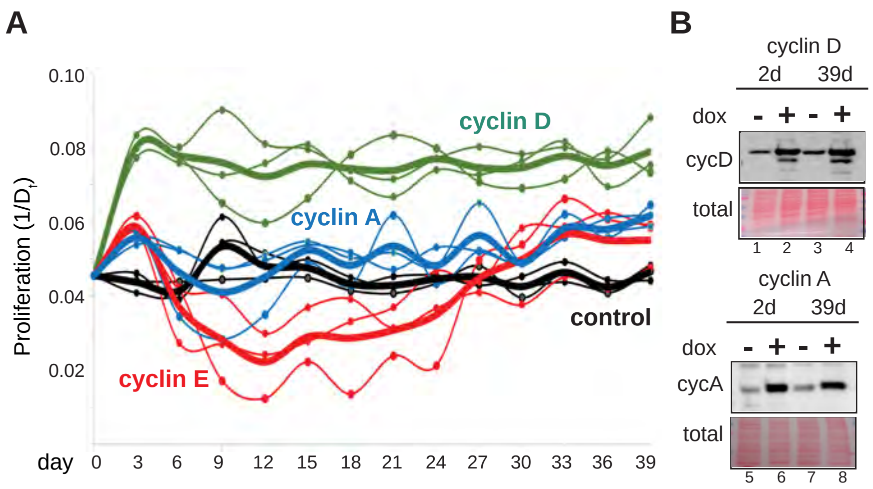


**Supplemental Figure 3. Adaptation to chronic cyclin D or cyclin A overproduction**. (A) Proliferation rate for uninduced control cells (black) and cells overproducing cyclin D1 (100 ng/ml, green) or cyclin A2 (100 ng/ml, blue) and cyclin E1 (20 ng/ml, red, from Figure 3A). (B) Immunoblot analysis of cyclin D1 (top) and cyclin A2 (bottom) after 2 or 39 days of proliferation under constant inducing conditions.

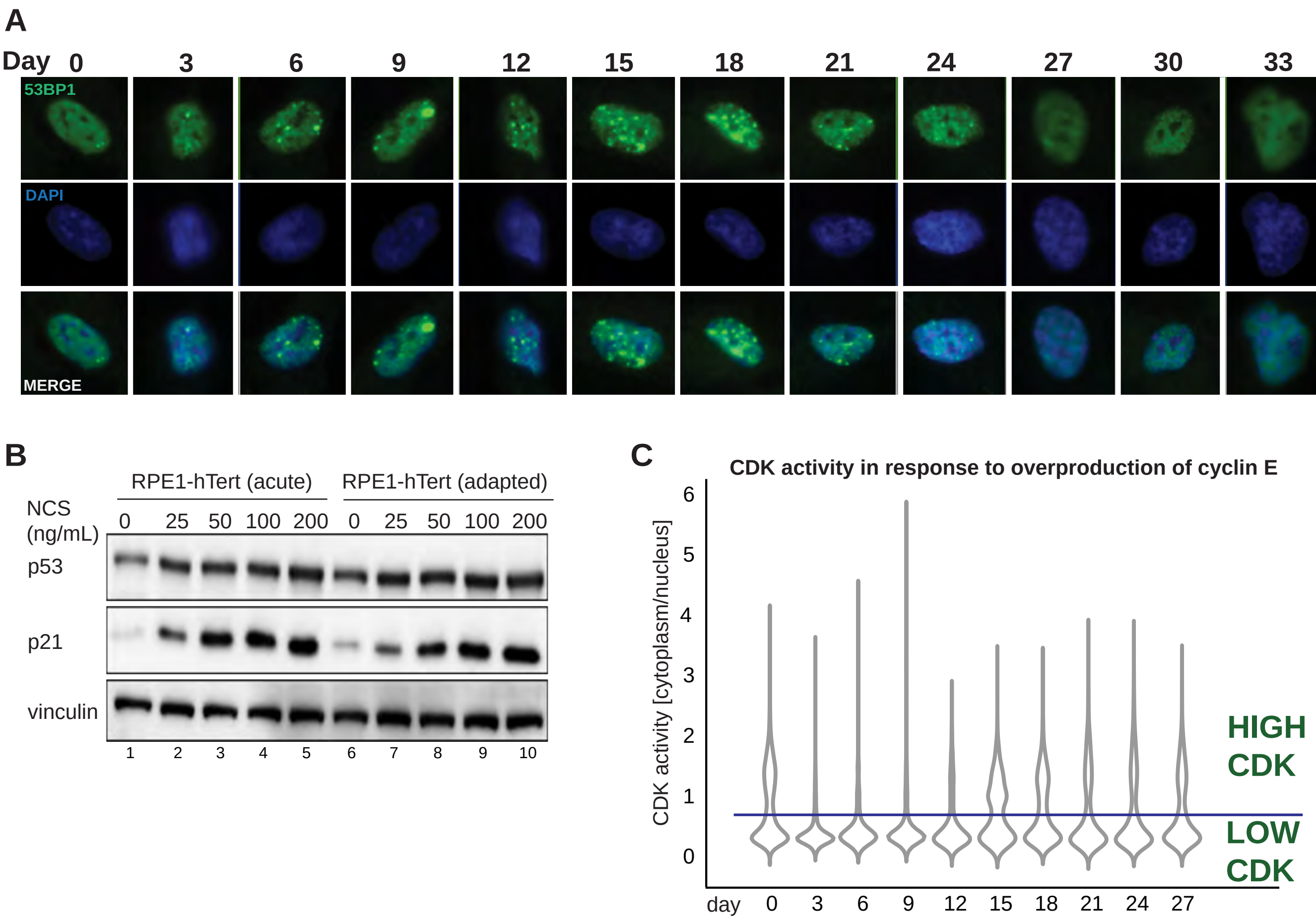


**Supplemental Figure 4. DNA damage markers and CDK activity during adaptation to cyclin E overproduction**. (A) Representative immunofluorescence images of cells fixed at three-day intervals during adaptation to cyclin E overproduction. DNA damage was scored as bright 53BP1 foci (green), and DNA was stained with DAPI (blue). (B) Cells were induced with 20 ng/mL doxycycline to overproduce cyclin E for 2 days or after adaptation (30+ days), then treated with the indicated concentrations of NCS for 48 hours. Endogenous p53, p21, and vinculin (as a loading control) in whole cell lysates were detected by immunoblotting. (C) CDK reporter activity at the end of each 72 hours of live imaging during the course of adaptation. The line at 0.75 marks a threshold of CDK reporter activity above which cells are committed to S phase, in S, or G2 phase.

S5

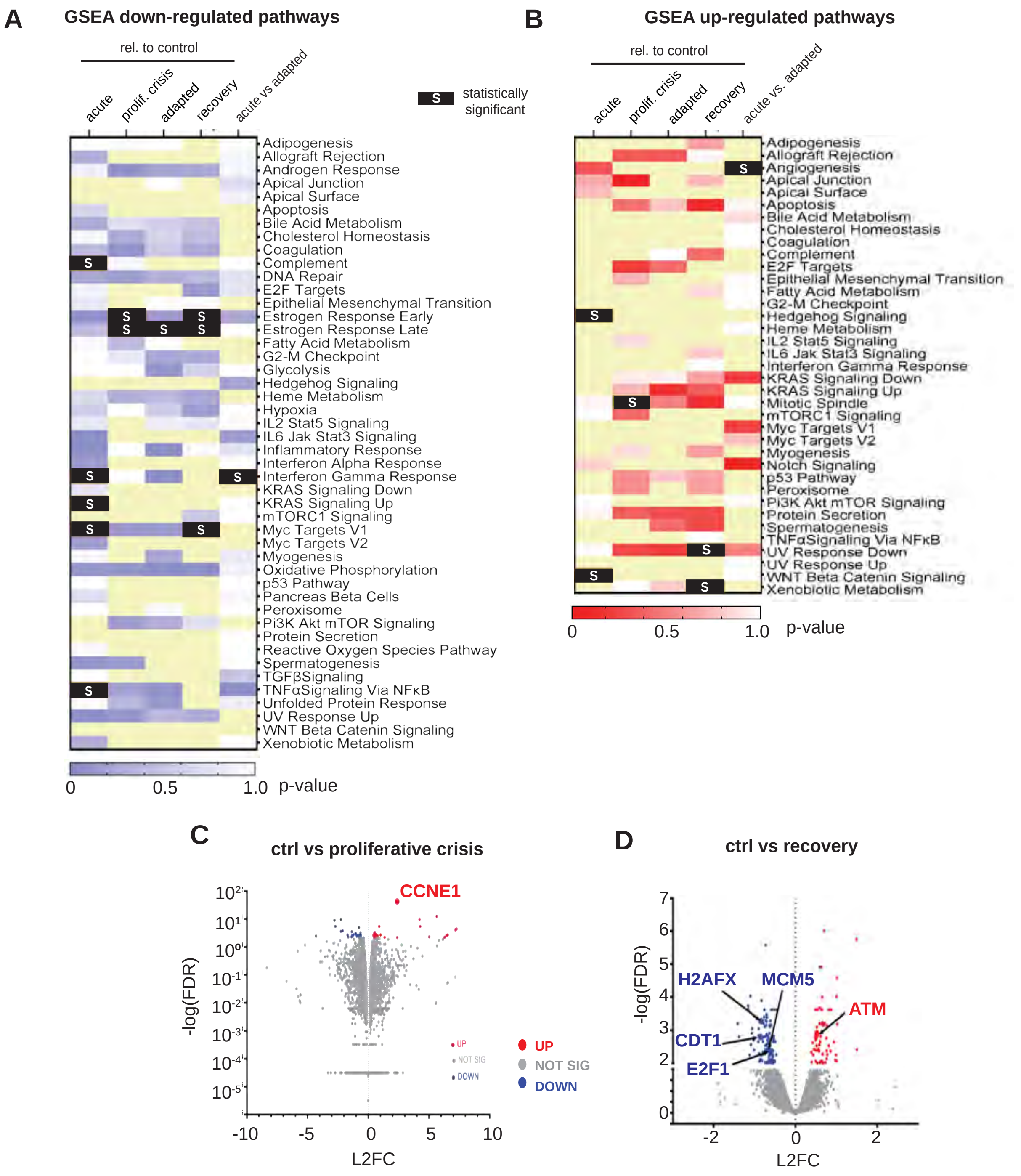


**Supplemental Figure 5. Individual gene and pathway changes during adaptation to cyclin E overproduction**. (A) Heat map of down-regulated pathways in the indicated pairwise comparisons using GSEA shaded by p-value. Statistically significant pathway changes defined by nominal p-value score <0.05 are highlighted in black with an “S” (light yellow boxes indicate no data) (B) Heat map of up-regulated pathways in the indicated pairwise comparisons using GSEA shaded by p-value. Statistically significant pathway changes are highlighted in black with an “S” (light yellow boxes indicate no data). (C) Volcano plot of all detected mRNAs comparing control to proliferative crisis (day 0 vs ~days 9-12) after induction; statistically significant upregulated or downregulated genes are marked in red or blue respectively. Gene expression changes below the threshold for significance are marked with gray dots. (D) As in C comparing control to recovered cells after doxycycline withdrawal; selected origin licensing and DNA damage response genes are highlighted. See also Supplemental Table 1.

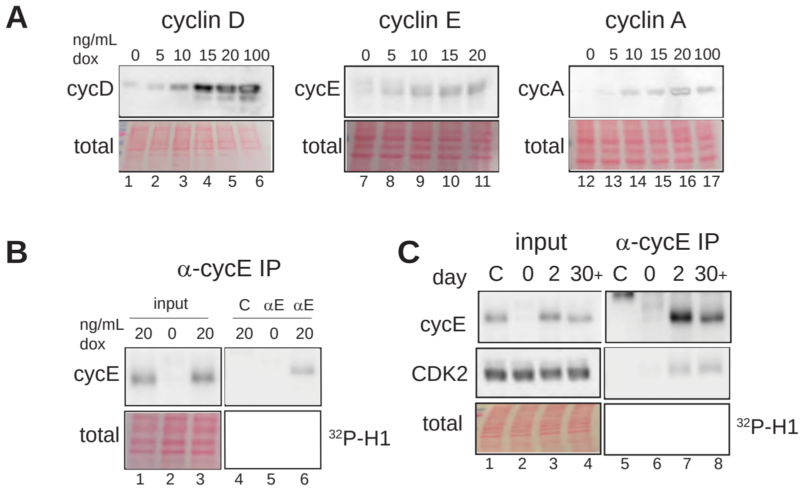


**Supplemental Figure 6. Lighter exposures of Figures 1A, 1D, and 3B.** (A) Lighter exposures of Figure 1A. (B) Lighter exposures of Figure 1D. (C) Lighter exposures of Figure 3B.
